# An Exhaustive Multiple Knockout Approach to Understanding Cell Wall Hydrolase Function in *Bacillus subtilis*

**DOI:** 10.1101/2021.02.18.431929

**Authors:** Sean A. Wilson, Raveen K. J. Tank, Jamie K. Hobbs, Simon J. Foster, Ethan C. Garner

## Abstract

Most bacteria are surrounded by their cell wall, containing a highly crosslinked protective envelope of peptidoglycan. To grow, bacteria must continuously remodel their wall, inserting new material and breaking old bonds. Bond cleavage is performed by cell wall hydrolases, allowing the wall to expand. Understanding the functions of individual hydrolases has been impeded by their redundancy: single knockouts usually present no phenotype. We used an exhaustive multiple-knockout approach to determine the minimal set of hydrolases required for growth in Bacillus subtilis. We identified 42 candidate hydrolases. Strikingly, we were able to remove all but two of these genes in a single strain; this “Δ40” strain shows only a mild reduction in growth rate, indicating that none of the 40 hydrolases are necessary for growth. The Δ40 strain does not detectably shed old wall, suggesting that turnover is not essential for growth. The remaining hydrolases in the Δ40 strain are LytE and CwlO, previously shown to be synthetically lethal. Either can be removed in Δ40, indicating that either hydrolase alone is sufficient for cell growth. Screening of environmental conditions and biochemistry revealed that LytE activity is inhibited by Mg2+ and that RlpA-like proteins may stimulate LytE activity. Together, these results suggest that the only essential function of cell wall hydrolases in B. subtilis is to enable cell growth by expanding the wall and that LytE or CwlO alone is sufficient for this function. These experiments introduce the Δ40 strain as a tool to study hydrolase activity and regulation in B. subtilis.

**IMPORTANCE:** In order to grow, bacterial cells must both create and break down their cell wall. The enzymes that are responsible for these processes are the target of some of our best antibiotics. Our understanding of the proteins that break down the wall – cell wall hydrolases – has been limited by redundancy among the large number of hydrolases many bacteria contain. To solve this problem, we identified 42 cell wall hydrolases in *Bacillus subtilis* and created a strain lacking 40 of them. We show that cells can survive using only a single cell wall hydrolase; this means that to understand the growth of *B. subtilis* in standard laboratory conditions, it is only necessary to study a very limited number of proteins, simplifying the problem substantially. We additionally show that the Δ40 strain is a research tool to characterize hydrolases, using it to identify 3 ‘helper’ hydrolases that act in certain stress conditions.

## INTRODUCTION

Most bacterial cells are surrounded by a peptidoglycan (PG) cell wall – a load-bearing structure that protects cells from lysing due to their high internal turgor (1). Bonds must be broken in the PG for cells to expand during growth (2). PG is built from disaccharide subunits linked to stem peptides. As new PG is inserted into the wall, the disaccharides are polymerized into long chains, and their stem peptides are crosslinked into the existing wall (3).

The enzymes that break PG bonds are termed cell wall hydrolases (hereafter ‘hydrolases’). Hydrolases fall into several broad categories with different chemical specificities (4). Amidases cleave the stem peptide from the sugar subunit. Endopeptidases cleave bonds between peptides within the stem peptide. Lytic transglycosylases (LTGs) and lysozymes (both of which are muramidases), cleave between the disaccharide subunits (MurNAc-GlcNAc), reversing the transglycosylase reaction that polymerizes glycan chains. Glucosaminidases target the other bond between sugar subunits (GlcNAc-MurNAc), reversing a cytoplasmic step of PG synthesis. The LTG reaction mechanism is not a hydrolysis reaction; however, to avoid introducing new terminology and to improve readability, we will use the term ‘hydrolases’ generally to refer to all PG cleavage enzymes, including LTGs. A wide array of different protein domains are capable of hydrolase activity – for example, there are at least 7 distinct domains with LTG activity and well over 100 distinct domains with hydrolase activity discovered thus far (5, 6).

Hydrolase activity is essential: without the breakage of PG bonds, the cell wall cannot expand to accommodate the accumulating biomass it contains (2). Hydrolases are also involved in a variety of other processes that require modification of the cell wall: turning over old PG, cell separation, and sporulation, conjugation, and motility (4, 7). Perhaps owing to the multiple cellular functions that require hydrolases, many bacteria have a large number of hydrolases. *Bacillus subtilis* and *Escherichia coli,* for example, each contain at least 20 hydrolases (4, 8). The large number of hydrolases in each bacterium, combined with a high degree of functional and enzymatic redundancy between them, has made it difficult to identify specific cellular functions for many hydrolases. Single knockouts rarely present clear phenotypes due to compensation by other hydrolases (4, 9). However, multiple-knockout approaches in *B. subtilis* have been successful in revealing the importance of LytE and CwlO for cell growth, uncovering the role of LytC and LytD in cell wall turnover, and identifying LytE, LytF, and CwlS as cell separation hydrolases (4, 10, 11).

*lytE* and *cwlO* had been previously shown to be synthetically lethal when both are deleted in *B. subtilis* (10, 12). The requirement of LytE or CwlO for cell growth was demonstrated via microscopy and genetics: upon depletion of LytE in a *cwlO* null mutant, or vice versa, cell elongation slows and then stops completely before cells lyse (12). To test whether any other hydrolases were essential for *B. subtilis* growth, we employed an exhaustive multiple-knockout approach. We created a minimal hydrolase strain that allows the study of hydrolases in isolation, making it easier to assign functions to uncharacterized hydrolases. Using this multiple hydrolase knockout strain, it is straightforward to assay the biochemical activity and determine the effect of hydrolases alone or in any desired combination on phenotypes like cell width, cell wall turnover, cell growth, or any other process.

## RESULTS

### Construction of a multiple hydrolase knockout strain

To identify the minimal set of hydrolases required for growth in *B. subtilis* PY79, we constructed a strain in which we sequentially removed as many hydrolases as possible. We used PHMMER to screen the *B. subtilis* proteome for proteins containing cell wall hydrolase domains present in known hydrolases (4, 5, 8, 13, Table S2). The results of this search are shown in Table 1. Cell wall hydrolases present in *B. subtilis* 168 but not present in PY79, our wild-type (WT) background, are included for completeness, though we did not generate knockouts for these. Candidate hydrolases previously shown to be unable to degrade intact PG (indicated in Table 1) were also not knocked out, nor were candidate hydrolases with transmembrane domains, as these are unlikely to be able to reach into the cell wall space far enough to directly participate in growth. We found 56 candidate hydrolases in the initial search (all shown in Table 1); of these, 50 were present in PY79. 4 were excluded due to previous work demonstrating lack of activity against intact PG (AmiE, NagZ, LytB, and PgdS), and 4 were excluded due to being membrane-bound (MltG, SweC, SpoIIQ, and SpoIVFA), leaving 42 candidate hydrolases for our study.

**Table 1:**
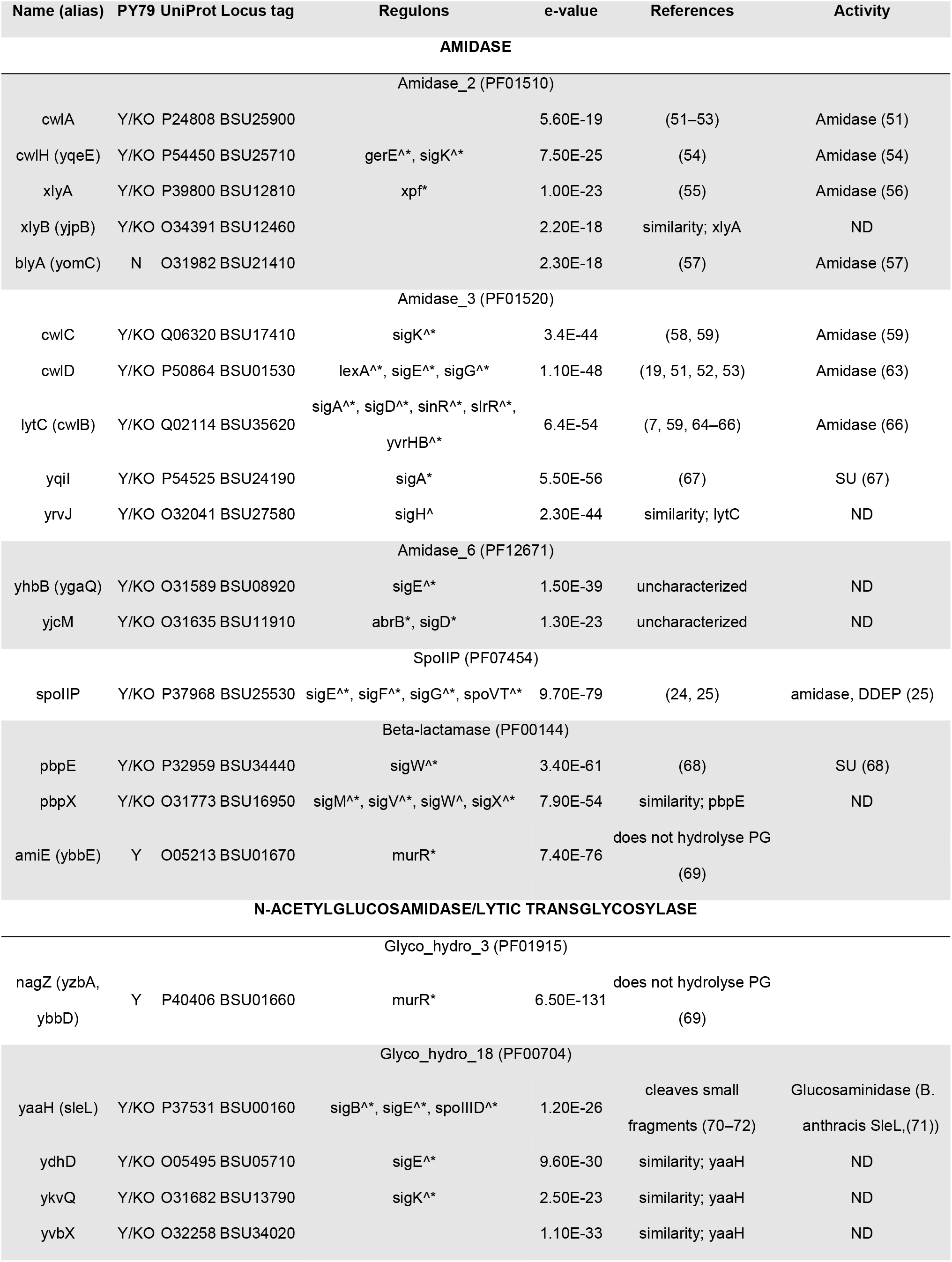
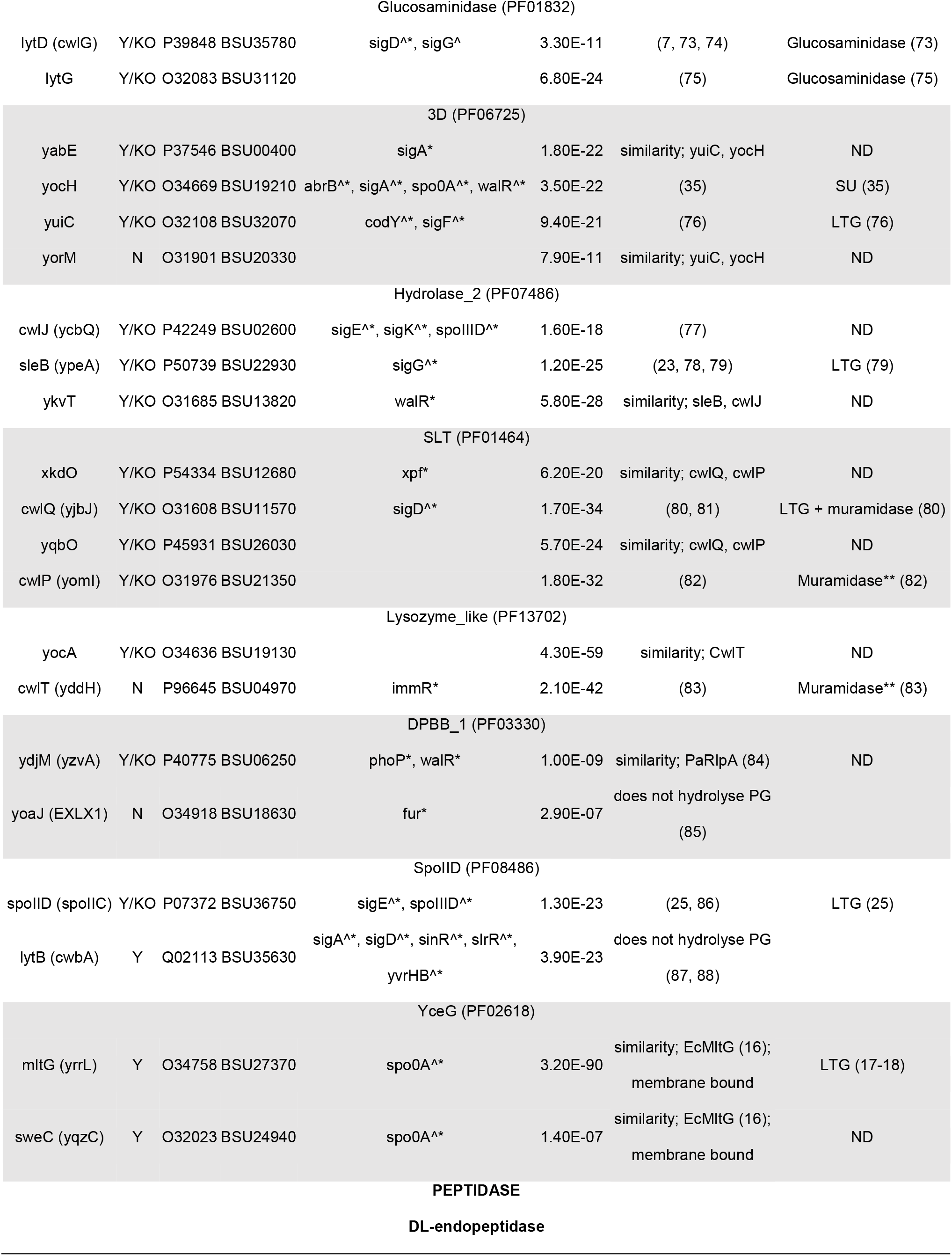
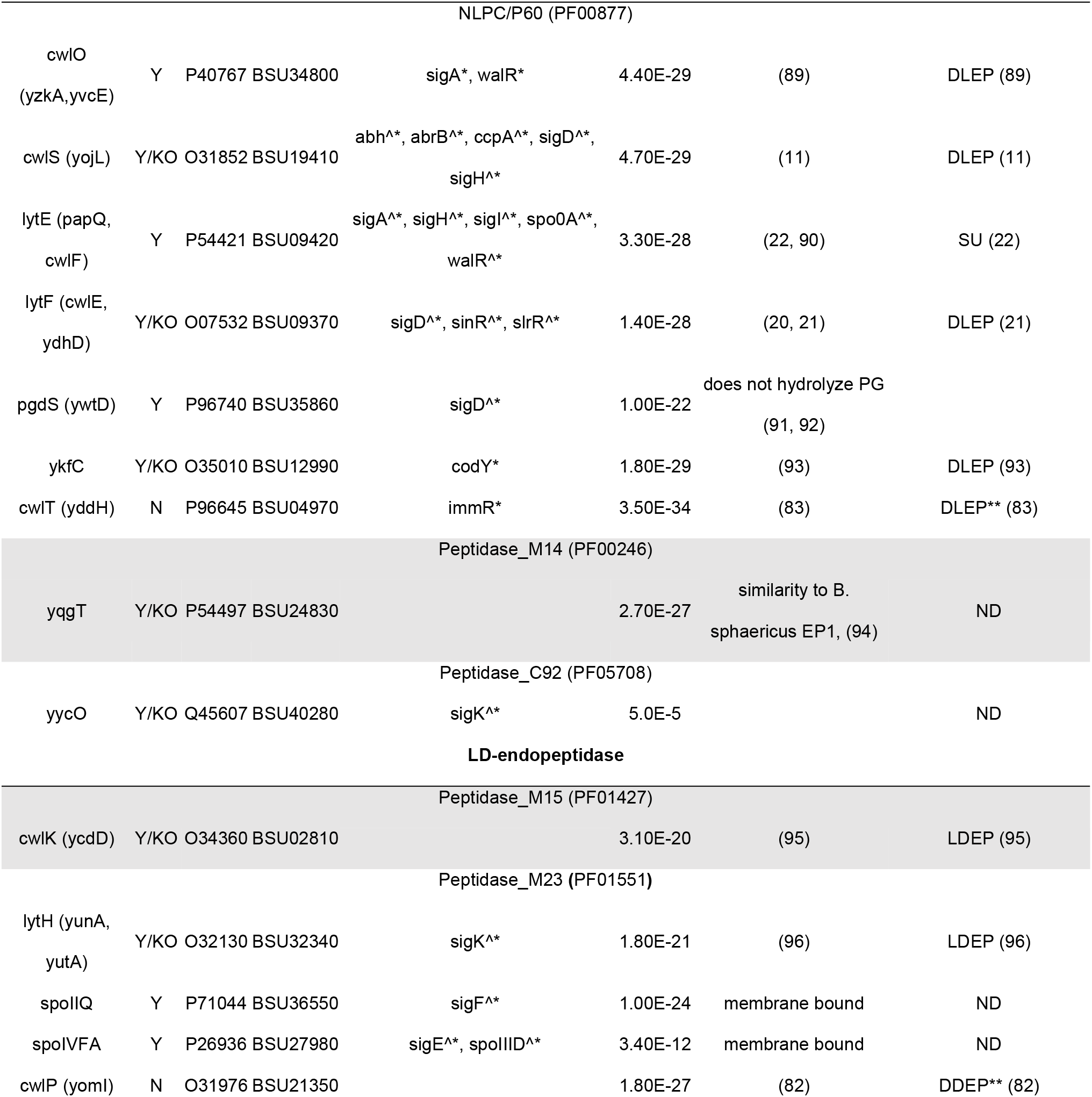
List of cell wall hydrolases in *Bacillus subtilis* identified using PHMMER. Cell wall hydrolases were identified via a PHMMR (13) search with default parameters of the *B. subtilis* subsp. 168 and *B. subtilis* subsp. PY79 proteome for PFAM domains associated with known cell wall hydrolases (Table S2). For each hydrolase, we report its name (and any aliases), whether it is knocked out in the Δ40 strain (KO) or present in PY79 (Y/N), its UniProt accession number, its locus tag, any reported regulons it is a member of (^ indicates source Faria et al. 2016, * indicates source SubtiWiki (97)), the PHMMR search significance e-value, and any relevant references showing its biochemical activity. Abbreviations: DLEP, D,L-endopeptidase; LDEP, L,D-endopeptidase; DDEP, D,D-endopeptidase; LTG, lytic transglycosylase; SU, specificity untested (only cell wall degradative activity shown). **CwlP and CwlT are two-domain cell wall hydrolases and so appear twice on this table. Only the activity for the specific PFAM domain is listed in the Activity column.

We next generated single knockouts for each of the candidate hydrolases by replacing the gene with an antibiotic resistance cassette flanked by loxP sites. We then sequentially combined all knockouts into a single strain, using Cre-lox mediated loop outs to remove markers when necessary (Figure 1). After each loop out step, we verified deletion of all modified loci by PCR. After all knockouts had been combined into a single strain, whole-genome sequencing was used to confirm all deletions and to identify any genomic rearrangements or mutations that occurred during the construction process. Despite the multiple rounds of transformation and loop outs this strain was subjected to, with up to 4 resistance cassettes removed simultaneously multiple times, we found no evidence of genomic rearrangements based on read coverage from DNA extracted during exponentially growing cells (Figure S1) (14), and only 8 SNPs leading to 5 point mutations in genes involved in unrelated processes (Table S1).

**Figure 1:**
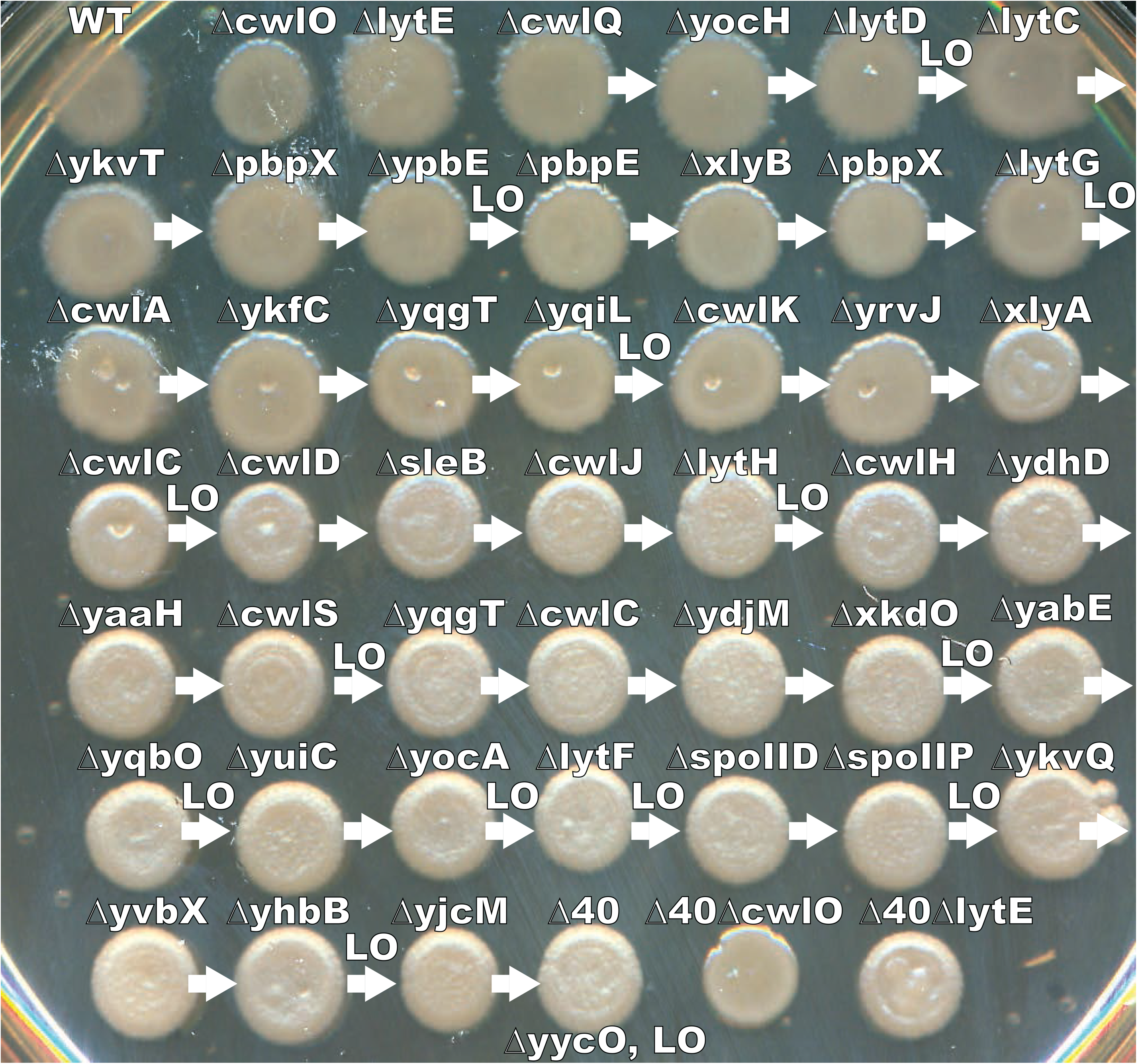
Construction of the Δ40 strain via sequential knockout and loopout. Colony morphology of each cloning intermediate for the Δ40 strain. WT cells were transformed with a series of resistance-cassette-marked knockouts (starting with *ΔcwlQ*). Periodically, antibiotic resistance cassettes were removed via Cre-loxP mediated loopout (indicated by LO). Arrows indicate sequential integrations (e.g., the strain indicated by *ΔyocH* contains *ΔyocH* and *ΔcwlQ*). Dense cell suspensions were spotted and incubated overnight to visualize colony morphology.

Ultimately, this effort produced a strain lacking 40 hydrolases, which we termed “Δ40”. The Δ40 strain is lacking all the identified hydrolases that met our criteria save two - LytE and CwlO, two synthetically lethal endopeptidases previously shown to be essential for growth (10). We were able to further knock out either *lytE* or *cwlO* in the Δ40 strain, but not both, due to their synthetic lethality.

### Hydrolase activity is greatly reduced in the Δ40 strain

To assess whether any other unidentified hydrolases remained in the Δ40 strain, we conducted PG profiling of both wild type (WT) cells and the Δ40 strain (15), allowing us to determine the abundance of hydrolase products in their cell walls (Figure 2, Table S3).

**Figure 2:**
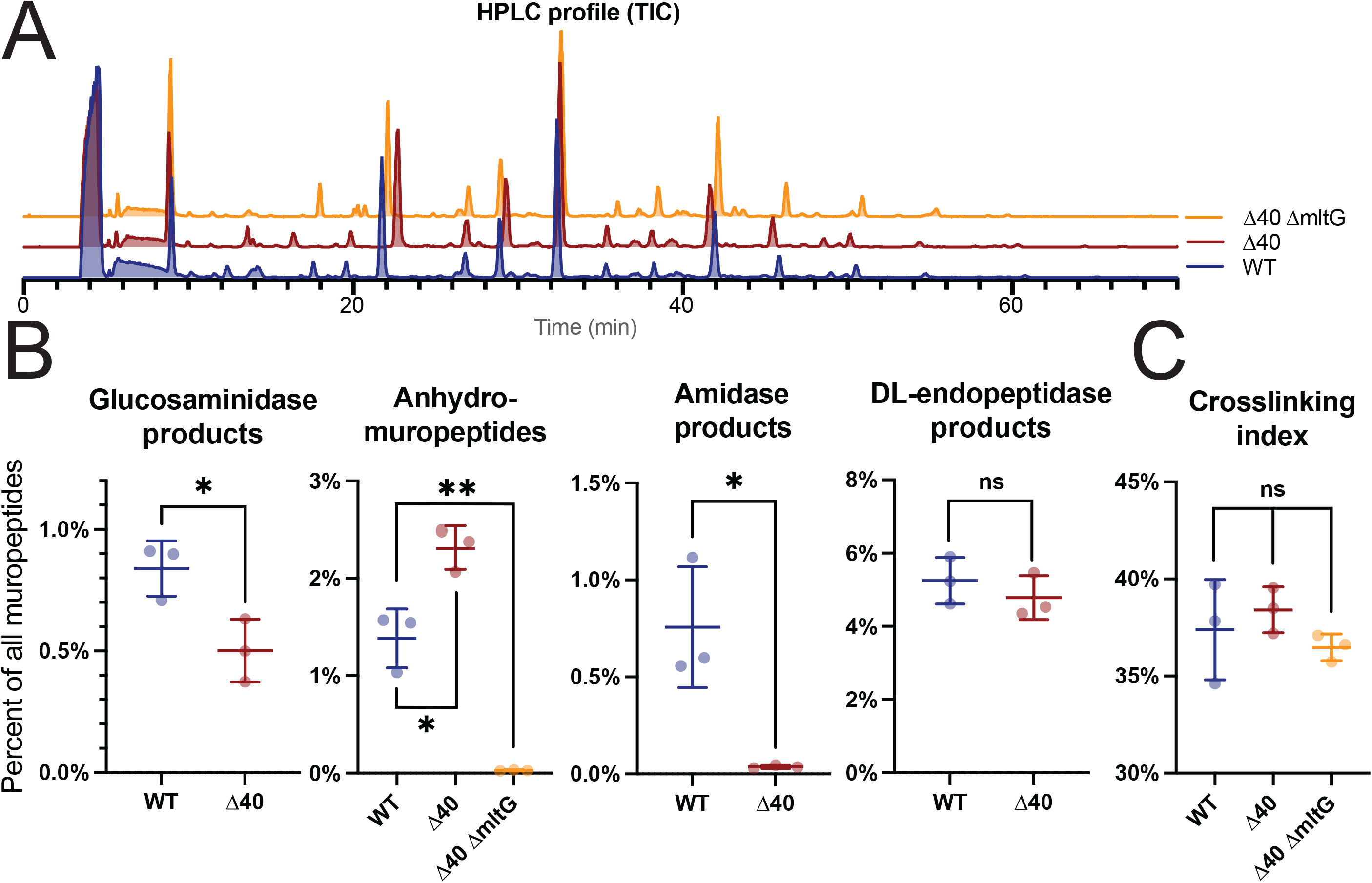
The Δ40 strain has a reduced cell wall hydrolytic complement. **A: HPLC analysis of isolated muropeptides in WT and Δ40 strains.** Purified cell walls were digested to yield soluble muropeptides which were separated and characterized via HPLC-MS. The total ion current (TIC) elution profile is shown. Strains used: PY79, WT; bSW431, Δ40; bSW537, Δ40 *ΔmltG*. **B: Identification of cell wall hydrolase products in WT and Δ40 cells by peptidoglycan profiling**. High-resolution mass spectrometry was used to identify separated muropeptides. Muropeptides missing a GlcNAc were classified as glucosaminidase products. Anhydromuropeptides were classified as LTG products. Crosslinked muropeptides lacking MurNAc-GlcNAc and MurNAc-GlcNAc itself was classified as amidase products. Crosslinked muropeptides lacking MurNAc-GlcNAc-L-Ala-iso-D-Glu, and MurNAc-GlcNAc-L-Ala-iso-D-Glu itself, was classified as D,L-endopeptidase products. For additional details about muropeptide classification, see Supplemental Table 2. Crosslinking index was calculated as in (98). For each set of hydrolase products, the sum of the MS intensity for those products was divided by the total MS intensity for all detected muropeptides (% of total). D,L-endopeptidase products are still present as expected, because the strain retains the D,L-endopeptidases LytE and CwlO. Amidase products are strongly reduced. Glucosaminidase products are reduced in abundance by roughly twofold. Lytic transglycosylase products are still present in the Δ40 strain but are strongly reduced if *mltG* is additionally knocked out. Strains used: PY79, WT; bSW431, Δ40; bSW537, Δ40 *ΔmltG*.

Our PG profiling assay has limitations: as PG profiling relies on muramidase digestion to yield soluble muropeptides for HPLC analysis, we could not use this assay to detect hydrolases with muramidase activity. Likewise, as D,D-endopeptidases produce products that are indistinguishable from unmodified PG, we cannot unambiguously assign specific PG products to D,D-endopeptidases in these experiments.

We compared the relative abundance of different PG hydrolase products in the Δ40 and WT strains (Figure 2B, Table S3). The Δ40 strain showed a very small amount of amidase activity (∼20-fold reduction vs WT, 0.8% vs 0.04% of all muropeptides, p=0.0161, unpaired t test) and a reduction of glucosaminidase activity (∼2-fold reduction vs WT, 0.8% vs 0.5% of all muropeptides, p=0.0279, unpaired t test), indicating that these classes of hydrolases had been successfully reduced in the Δ40 strain. The residual glucosaminidase activity could represent A) a yet unknown minor glucosaminidase with a novel fold or B) sample degradation during PG purification. We observed no change in D,L-endopeptidase activity in the Δ40 strain (5.2% vs 4.8% of all muropeptides, p=0.4094, unpaired t test), an expected result given Δ40 retains the D,L-endopeptidases LytE and CwlO. In agreement with previous work (15), L,D-endopeptidase activity was not detected in any strain. D,L-endopeptidases cleave between the mDap and iso-D-Glu residues in the stem peptide, while L,D-endopeptidases cleave between iso-D-Glu and L-Ala.

Unexpectedly, the Δ40 strain also showed an increase in LTG activity (Figure 2B, ∼1.75-fold increase vs WT, 1.4% vs 2.3% of all muropeptides, p=0.0125, unpaired t test). We found that this remaining LTG activity required MltG. Removing *mltG* from the Δ40 strain substantially reduced apparent LTG activity (Figure 2B, ∼80-fold reduction vs Δ40, 2.3% vs 0.03% of all muropeptides, p<0.0001, unpaired t test). MltG’s catalytic domain is predicted to be extracellular, although MltG is likely too small to reach far enough into the cell wall space to directly participate in cell wall expansion. MltG has been shown to be involved in membrane-proximal PG metabolism, cleaving PG at a specific distance from the membrane to produce 7-dissacharide long glycan strands (16–18).

Additionally, we measured the rate of autolysis in the Δ40 strain. Autolysis occurs when hydrolases become dysregulated and degrade the wall in an uncontrolled way, leading to cell lysis. Disruption of the energized membrane via energetic poisons, or treatment with antibiotics can cause autolysis (19). In *B. subtilis*, the hydrolases LytC and LytD are the main effectors of autolysis, with LytE and LytF additionally having smaller effects (7, 20).

*ΔcwlO* and *ΔlytE* cells had approximately WT rates of autolysis, with nearly 100% of cells being lysed after 5H of treatment with 75 mM sodium azide or 100 µg/ml ampicillin. In contrast, the Δ40 strain had a significantly slower rate of autolysis in both treatment conditions, with the Δ40 *ΔlytE* strain showing only a ∼10% reduction in OD_600_ after 24H of treatment and the Δ40 strain itself showing a ∼40% OD_600_ reduction after 24H. The nonzero autolysis rate in the Δ40 *ΔlytE* background could imply the involvement of CwlO in autolysis, could represent non-hydrolase-mediated lysis, or could suggest a remaining hydrolase with a minor role in autolysis.

### Cell growth and morphology are similar in the Δ40 strain relative to wild type

We next characterized the growth rate of the Δ40 strain. The Δ40 strain grew slightly slower than WT cells in rich, undefined media (LB), but grew at the same rate as WT cells in both rich, defined media (CH) and fully synthetic media (S7_50_ with glucose and amino acids, see Methods for details) (Figure 3A). This suggests that the activity of LytE and CwlO together are mostly sufficient for normal cell growth, although when pushed towards higher growth rates other cell wall hydrolases may contribute to growth. To investigate the individual effects of LytE and CwlO on the cell growth rate, we made knockouts of *lytE* and *cwlO* in both WT and Δ40 backgrounds. Δ40 Δ*lytE* and Δ40 Δ*cwlO* both exhibited a reduction in growth rate compared to Δ40 which was especially pronounced in LB media. We observed cell lysis in both Δ40 Δ*lytE* and Δ40 Δ*cwlO* strains in phase-contrast images, which could contribute to their slower growth rates as measured in bulk by OD_600_ (Figure 4A). On the other hand, Δ*lytE* or Δ*cwlO* in a WT background had the same growth rate as WT. This suggests either that in WT cells, other hydrolases participate in but are not strictly required for growth, or that LytE and CwlO are not being expressed highly enough to maintain normal growth on their own in the Δ40 background.

**Figure 3:**
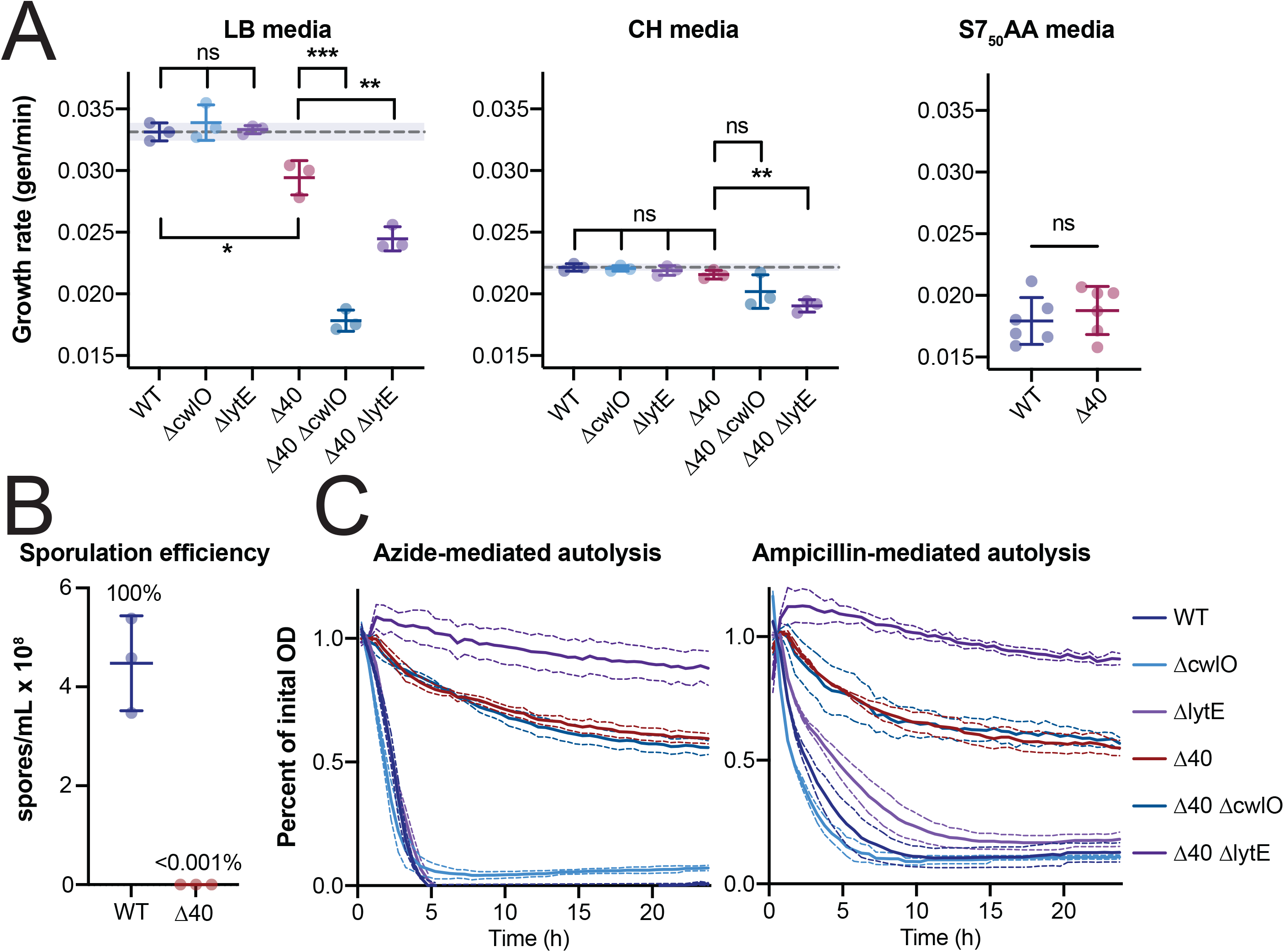
The Δ40 strain grows at a similar rate as WT cells, does not sporulate, and has a much slower autolysis rate in response to both sodium azide and ampicillin. **A (left): The Δ40 strain has a slower growth rate vs WT in LB media.** Cultures were grown in LB media at 37°C to an OD_600_ of ∼0.3-0.5, diluted to an OD_600_ of 0.05, and samples were collected every 6 minutes for 1 hrs (∼3 doublings). OD_600_ vs time plots were fit to a single exponential to obtain the growth rate. Each point represents the doubling time from a single experiment, and solid lines show mean and standard deviation. The dotted line shows the mean WT growth rate, for comparison. Δ40 has a slower growth rate to WT. *lytE* and *cwlO* knockouts grow much more slowly in the Δ40 background than in a WT background. Strains used: PY79, WT; bSW23, Δ*cwlO*; bSW295, Δ*lytE*; bSW431, Δ40; bSW433, Δ40 Δ*cwlO*; bSW435, Δ40 Δ*lytE*. **A (middle): The Δ40 strain has a similar growth rate to WT in CH media.** Cultures were grown in CH media at 37°C. Samples were collected and data was analyzed as in (A). While Δ40 has a similar growth rate to WT, *lytE* and *cwlO* knockouts grow more slowly in this background than in a WT background. Strains used: PY79, WT; bSW23, Δ*cwlO*; bSW295, Δ*lytE*; bSW431, Δ40; bSW433, Δ40 Δ*cwlO*; bSW435, Δ40 Δ*lytE*. **A (right): The Δ40 strain has a similar growth rate to WT in minimal media.** Cultures were grown in S7_50_AA media at 37°C. Samples were collected and data was analyzed as in (A). Strains used: PY79, WT; bSW431, Δ40. **B: The Δ40 strain is unable to sporulate.** Sporulation was induced by resuspension and sporulation efficiency was determined as in (24). The WT strain produced ∼10^8^ spores/mL while the Δ40 strain produced ∼100 spores/mL, all of which lacked the distinctive Δ40 colony morphology and likely represent contamination. Strains used: PY79, WT; bSW431, Δ40. **C: The Δ40 strain shows slower autolysis in response to sodium azide (left) and ampicillin (right).** Cultures were grown in CH media at 37°C to an OD_600_ of 0.5 (azide treatment) or 0.15 (ampicillin treatement) in baffled flasks with vigorous shaking, then diluted to an OD_600_ of 0.025 in a 150 µL of prewarmed CH in a 96-well plate. Sodium azide (75 mM) or ampicillin (100 µg/mL) was added and OD_600_ readings were taken using a plate reader every 2 minutes for 24H. The plate was shaken vigorously in between measurements. The Δ40 strain takes around 10 times as long to reach half the intial OD_600_ as the WT strain, and the Δ40 ΔlytE strain in particular has a strongly reduced rate of autolysis. Strains used: PY79, WT; bSW23, Δ*cwlO*; bSW295, Δ*lytE*; bSW431, Δ40; bSW433, Δ40 Δ*cwlO*; bSW435, Δ40 Δ*lytE*.

**Figure 4:**
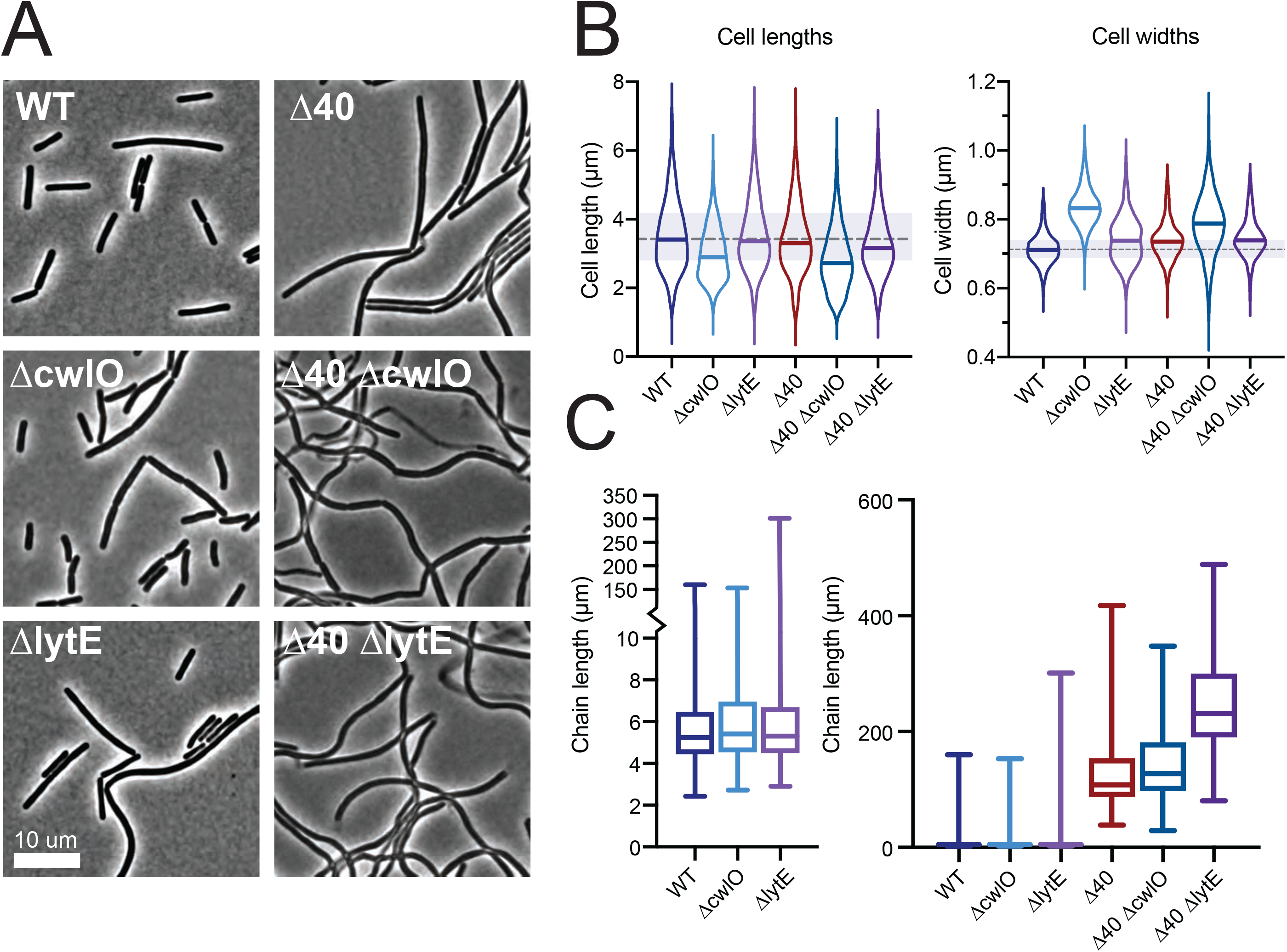
The Δ40 strain has mild shape defects but a significant chaining phenotype. **A: Representative phase contrast images of hydrolase mutant strains.** Δ40 cells primarily form long chains, Δ40 Δ*cwlO* cells have variable widths, and Δ40 Δ*lytE* cells sometimes have phase-light, lysed cells still attached to their poles (see Figure S2 for TEM images). Both Δ40 *ΔcwlO* and Δ40 *ΔlytE* have a population of phase-light, lysed cells. Scale bar is 10 µm. **B: Cell lengths (left) and widths (right) in hydrolase mutants.** Cells were labeled with membrane stain and imaged by epifluorescence microscopy. Cell dimensions were measured from these images using Morphometrics. Solid lines in violins show medians. Dashed line outside violins shows WT median for comparison. Shaded region outside violins shows WT quartiles. Strains used: PY79, WT; bSW23, Δ*cwlO*; bSW295, Δ*lytE*; bSW431, Δ40; bSW433, Δ40 Δ*cwlO*; bSW435, Δ40 Δ*lytE*. **C: Chain lengths in hydrolase mutants (left, zoomed view of right).** Chain length was measured manually from tiled and stitched phase contrast images. The Δ40 strain’s median chain length is around 30 times longer than the WT median chain length. Strains used: PY79, WT; bSW23, Δ*cwlO*; bSW295, Δ*lytE*; bSW431, Δ40; bSW433, Δ40 Δ*cwlO*; bSW435, Δ40 Δ*lytE*.

Next, we quantified cell dimensions in these strains using FM 5-95 membrane stain. Δ40 cells had a WT cell length and were 3% wider (Figure 4B, p<0.0001, unpaired t-test with Welch’s correction). Δ*cwlO* cells were 13% wider and 18% shorter than WT cells, a phenotype that persisted in the Δ40 Δ*cwlO* strain (Figure 4B, p<0.0001 for all comparisons: unpaired t-test with Welch’s correction for width comparisons, Mann-Whitney test for length comparisons). Δ40 Δ*cwlO* cells were less able to control their width as compared to Δ40 cells, having a 1.5x wider cell width distribution (Figure 4B, 7.5% vs. 11.33% coefficient of variation, F test p<0.001). In contrast, Δ*lytE* cells were only slightly wider than WT cells (Figure 4B, 1%, p<0.0001, unpaired t-test with Welch’s correction), and Δ40 Δ*lytE* cells were slightly narrower (Figure 4B, 1%, p<0.0001, unpaired t-test with Welch’s correction) than Δ40 strain alone, with a slight decrease in length (Figure 4B, p<0.0001, Mann-Whitney test). Thus, CwlO appears to be involved in cell width maintenance, as removing *cwlO* causes changes in cell width both in Δ40 and WT backgrounds, consistent with previous reports (12). Furthermore, given that removing *cwlO* increases the cell width coefficient of variation in the Δ40 background but does not increase the width variation when deleted from WT cells, other hydrolases must also have a role in width homeostasis.

We then quantified the chain length for the Δ40 strain and derivatives. Individual B. subtilis cells are often found connected to their siblings via a common cell wall septum as cell separation and cell division do not always occur at the same time in this organism. Cell separation requires the action of hydrolases that cleave between the two connected cells – several different hydrolases serve this purpose in B. subtilis, primarily LytF, CwlS, and LytE (11, 21, 22). In WT cells, the average chain length was comparable to the length of individual cells (Figure 4C, ∼4.5 µm per chain vs ∼3.5 µm per cell). The maximum chain length observed was 150 µm. *ΔcwlO* and *ΔlytE* mutants had a similar average chain length although the *ΔlytE* mutant had a larger maximum chain length (Figure 4C, 300 µm), consistent with the known role of LytE in cell separation.

In contrast, the Δ40 strain had a significant increase in the average chain length (Figure 4C, 120 µm), nearly as long as the longest observed WT chain. The Δ40 *ΔcwlO* strain was similar to the Δ40 strain, while the Δ40 *ΔlytE* strain had a large increase in the average chain length (250 µm), almost double that of the longest observed WT chain. Because the Δ40 *ΔlytE* strain lacks all cell separation hydrolases, the remaining cell separation in this strain was likely due to mechanical tearing of cells under the vigorous shaking conditions needed to be able to measure accurate culture OD_600_ for these experiments; the ends of the chains had visible, phase-light debris resembling torn cells still attached visible by TEM (Figure S3). In gentler culture conditions on a roller drum, this strain grows as a large clump of cells visible to the naked eye.

Additionally, we tested the ability of the Δ40 strain to sporulate. Hydrolases are involved in both entry into sporulation, as well as exit from the spore during germination (23–25). We found that the Δ40 was not able to sporulate (Figure 2B, p=0.3859, one sample t test vs efficiency of 0), likely because it lacks SpoIID and SpoIIP, causing a block at the engulfment stage of sporulation (25).

### Δ40 cells do not detectably turn over their cell wall

Hydrolases are involved in cell wall turnover, where old PG material is shed from the cell wall (26). We measured the rate of cell wall turnover of both WT and Δ40 cells using pulse-chase labeling with the radioactive cell wall precursor ^3^H-N-acetylglucosamine (^3^H-GlcNAc). This revealed that, while WT cells turn over PG at a rate of about 50% per generation in agreement with previous work (26), turnover in Δ40 strain was not detectable, with a rate not significantly different from zero (Figure 5A, p=0.4837, one sample t test vs. rate of 0). These results suggests that LytE and CwlO, the only identifiable remaining hydrolases in the Δ40 strain, likely do not contribute to cell wall turnover. Furthermore, this data suggests that cell wall turnover is not an essential process: cell growth only requires the cleavage of bonds so the cell can expand.

**Figure 5:**
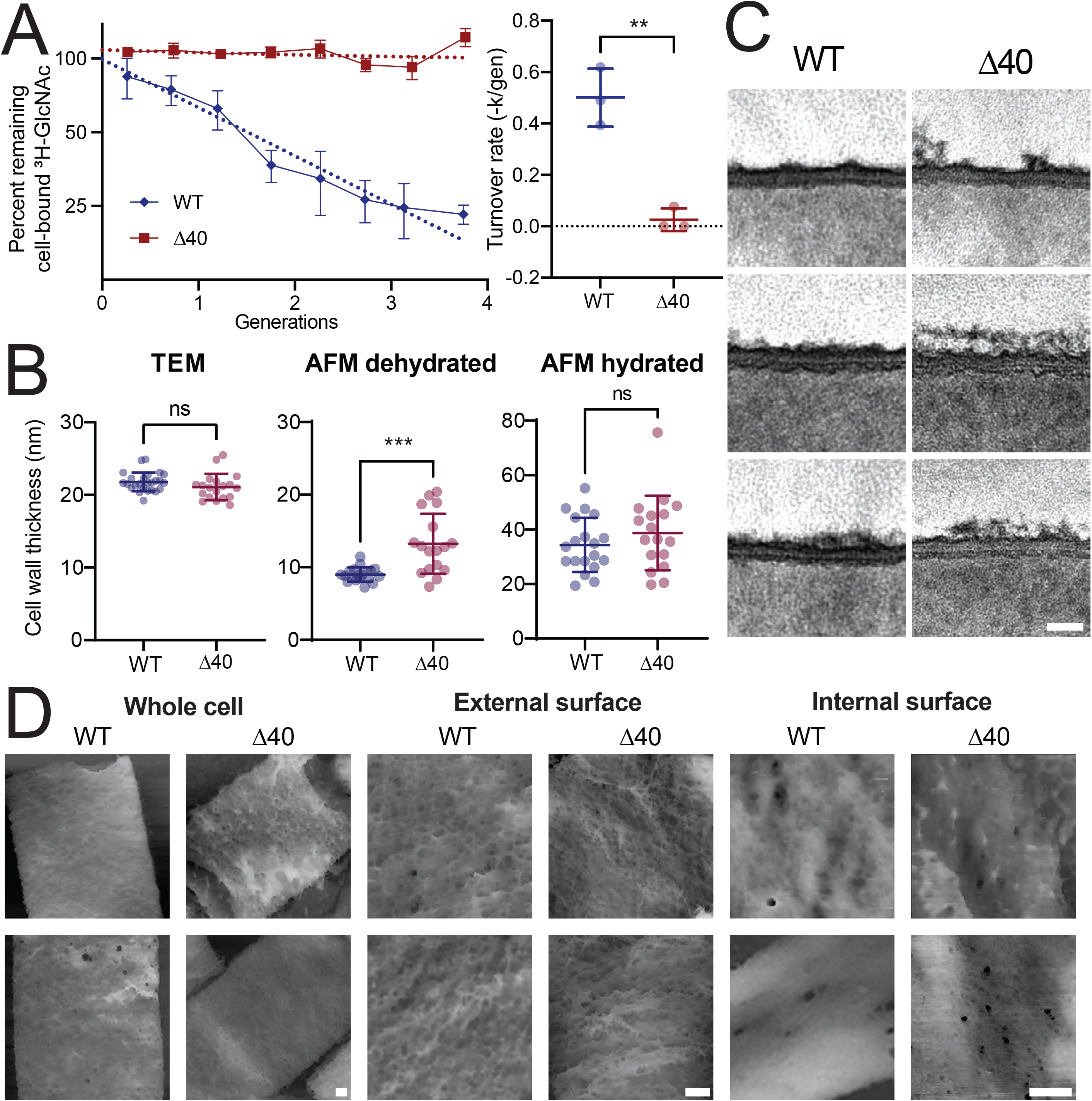
The Δ40 strain does not detectably turn over cell wall. **A: Cell wall turnover rate is negligible in the Δ40 strain.** *Left*: Pulse-chase radiolabel measurements were used to determine the cell wall turnover rate. Cells were labeled with H^3^-GlcNAc, which incorporates into the cell wall. The ^3^H-GlcNAc was then washed out and radioactivity was subsequently measured for 3 generations. A decrease in radioactivity indicates that material is being removed from the cell wall, e.g. that cell wall is turning over. Each experiment was replicated at least 3 times. Dotted lines show single exponential fit to mean data. *Right*: Single exponential fits to each experiment at left. Each point represents the time constant (-k) obtained from a fit to a single experiment. Error bars show SD. The Δ40 turnover rate is not significantly different from zero (one sample t test, p=0.4837). Mean initial radioactivity was 167766 DPM/OD_600_ for the WT strain and 174334 DPM/OD_600_ for the Δ40 strain. Strains used: PY79, WT; bSW431, Δ40. **B: Cell wall thickness in the Δ40 strain.** Cell wall thickness was measured via transmission electron microscopy and atomic force microscopy as described in Methods. Briefly, for TEM imaging, exponentially growing cells were fixed, osmicated, stained with uranyl acetate, embedded in Embed 812, sectioned, and imaged without additional staining. For AFM imaging, exponentially growing cells were boiled, broken, protease treated, and then adhered to mica for imaging. Each point is the mean cell wall thickness measured for a single cell. WT AFM measurements are reproduced from (99), Δ40 AFM measurements were performed as part of this study. Error bars show SD. Strains used: PY79, WT (for TEM measurements); 168, WT (for AFM measurements); bSW431, Δ40. **C: Representative TEM images of cell wall thickness.** Representative TEM images of cell wall thickness analyzed in B. Strains used: PY79, WT; bSW431, Δ40. Scale bar is 20 nm. **D: Representative AFM images of sacculi.** Representative AFM images of cell walls analyzed in B. Strains used: 168, WT; bSW431, Δ40. Scale bars are 100 nm.

As hydrolase-deficient mutants have been shown to have altered cell wall thickness (27, 28) we measured the cell wall thickness of the Δ40 strain using transmission electron microscopy (TEM) and atomic force microscopy (AFM). We found that the wall was significantly thicker only in dehydrated samples measured using AFM (Figure 5B, center, p=0.0007, unpaired t test); measurements on TEM images or on hydrated AFM samples showed no significant differences (TEM: Figure 5B, left, p=0.1382, unpaired t test; hydrated AFM: Figure 5B, right, p=0.2887, unpaired t test). The WT cell wall was more uniform in appearance in both TEM and AFM images, while the Δ40 strain had more heterogeneity in density and thickness, in particular on the outer face of the wall, with an increase in the presence of “ruffles” on the outer face of the cell wall in the Δ40 strain (Figure 5C,D). These “ruffles” may represent the additional old cell wall material present due to the strongly reduced turnover rate. The internal face of the cell wall appeared denser than the external face of the wall in both the WT and Δ40 strains (Figure 5D). The internal face of the cell wall in the Δ40 strain appeared to have both a denser meshwork and an increased number of larger pores as compared with the WT strain.

Substantial changes to the cell wall ultrastructure occur during sample prep for TEM, especially in the outer layers of the cell wall (29–31); it would be interesting to apply additional, less perturbative EM modalities such as cryo-electron microscopy to the Δ40 strain to help clarify the exact nature of the changes to the Δ40 cell wall. It is possible that both TEM and hydrated AFM highlights mostly the denser, newer cell wall material, while dehydrated AFM allows visualization of all the cell wall, including the more loosely-bound older wall material.

### Δ40 Δ*cwlO* cells are sensitive to various stresses, including ionic stress

Although the Δ40 strain grew mostly normally under our standard lab conditions, we wondered whether the absence of so many hydrolases would sensitize cells to stress conditions. We used a spot dilution assay to measure the viability of our strains under a variety of stress conditions: temperature, ionic stress, pH, and osmotic stress (Figure 6). In all conditions, including our control (37°C), Δ40 cells had fewer CFUs than WT. This is expected because Δ40 cells grow in long chains, and thus cells cannot readily separate into individual CFUs. In all stress conditions Δ40 cells were similarly viable to WT cells, as were Δ*lytE,* Δ*cwlO,* and Δ40 Δ*lytE* cells. However, Δ40 Δ*cwlO* cells were susceptible to multiple stresses, including low pH, low temperature, and ionic stress.

**Figure 6:**
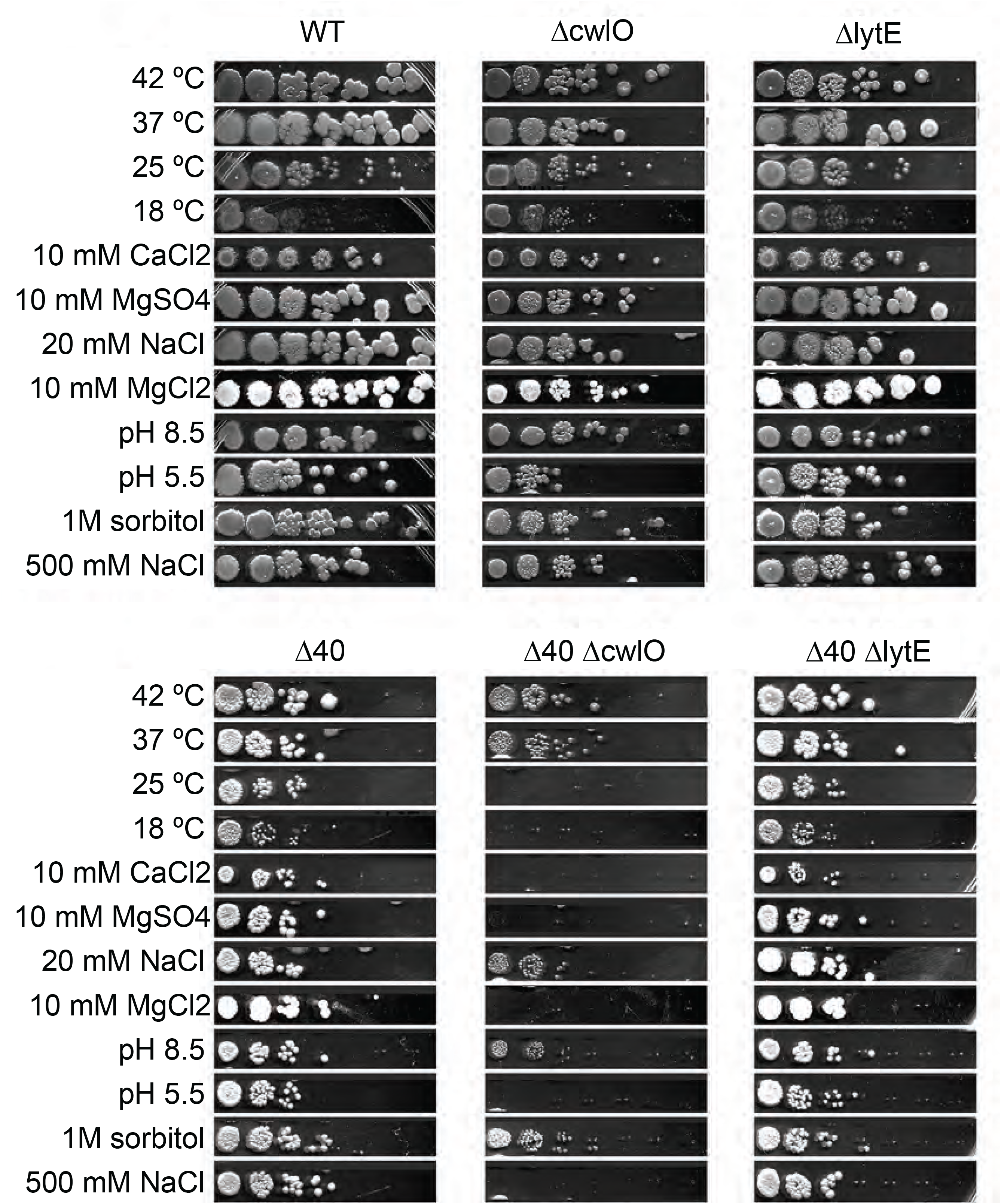
The Δ40 strain has similar viability to WT in a range of stress conditions, but Δ40 Δ*cwlO* is sensitive to ionic, cold, and low pH stress. Spot dilution assays of different strains under various stress conditions. Cultures of each strain were plated in a 1:10 dilution series onto LB plates containing various stressors and grown overnight at the specified temperature, or at 37°C if not indicated. Most conditions supported normal growth, but growth of the Δ40 Δ*cwlO* strain was inhibited at 25°C, pH 5.5, or with the addition of 10 mM MgCl_2_, 10 mM MgSO_4_, 10 mM CaCl_2_, or 300 mM NaCl. Strains used: PY79, WT; bSW23, Δ*cwlO*; bSW295, Δ*lytE*; bSW431, Δ40; bSW433, Δ40 Δ*cwlO*; bSW435, Δ40 Δ*lytE*.

We were particularly intrigued by the susceptibility of Δ40 Δ*cwlO* to Mg^2+^. Mg^2+^ is coordinated between PG and teichoic acids (32), and this Mg^2+^ binding is thought to give structural stability to the cell wall (31, 33). High levels of Mg^2+^ are often protective against cell wall perturbations, including knockouts of hydrolases, PBPs, or components of the Rod complex (10, 34); thus, the Mg^2+^ sensitivity of the Δ40 Δ*cwlO* strain seemed counterintuitive. Our experiments indicated Δ40 Δ*cwlO* cells were sensitive to both Ca^2+^ and Mg^2+^; growth was inhibited by the addition of 10 mM MgCl_2_, 10 mM MgSO_4_, and 10 mM CaCl_2_, but not by the addition of 20 mM NaCl, suggesting that the growth inhibition was not due to changes in ionic strength or chloride ions. We did observe growth inhibition due to ionic stress at far higher salt concentrations (500 mM NaCl). Notably, cells were not sensitive to an equivalent osmotic stress (1M sorbitol), indicating the sensitivity is to ionic stress, not osmotic stress.

As Δ*cwlO* mutants in the WT background were Mg^2+^ insensitive, we sought to identify which hydrolases caused cells to be sensitive to Mg^2+^ when they were removed. To find these hydrolases, we returned to intermediate strains used to construct the Δ40 strain, which are missing subsets of hydrolases. We transformed a *cwlO* knockout into these intermediate strains, then screened these crosses for the same small colony phenotype and the Mg^2+^ sensitivity seen in the Δ40 Δ*cwlO* strain. This identified two genes: *yabE* and *ydjM*. Notably, during construction of the Δ40 strain, we had noticed that *yocH* seemed significant – at several intermediate verification steps, a WT copy of *yocH* had reintegrated itself during transformation with genomic DNA from single KO strains – we therefore used PCR product for all transformations after this. Furthermore, a *ΔydjM ΔyocH ΔcwlO* mutant was previously demonstrated to be sick, with short and sometimes anucleate cells (10). Because *yabE*, *ydjM*, and *yocH* have similar hydrolase domains, and because *yocH* and *ydjM* had been identified previously to be involved in a synthetic sick interaction with *cwlO,* we additionally tested whether the removal of *yocH* contributed to the Δ40 *ΔcwlO* Mg^2+^ sensitivity phenotype, and found that it did.

In total, we identified three genes, *yabE*, *ydjM*, and *yocH*, whose absence in a Δ*cwlO* background caused the Mg^2+^ sensitivity: A *ΔyabE ΔydjM ΔyocH ΔcwlO* strain showed a similar stress profile to Δ40 Δ*cwlO*, including sensitivity to MgCl_2_ and CaCl_2_ (Figure 6A). *yabE*, *ydjM*, and *yocH* are 3 uncharacterized RlpA-like superfamily domain-containing proteins expressed during exponential growth. Like *lytE* and *cwlO, yocH* and *ydjM* are in the *walR* regulon, while *yabE* is regulated by *sigA* (Table 1). All are likely lytic transglycosylases: *yocH* has been shown to have lytic activity and has homology to the *E. coli* lytic transglycosylase *mltA* (35), and all three share a similar catalytic domain. Because *yabE*, *ydjM*, and *yocH* all contain a RlpA-like protein domain, we refer to these genes collectively as RLPAs, and to the triple deletion of all three genes as ΔRLPAs.

### LytE is inhibited by Mg^2+^ *in vitro* and *in vivo*, and RLPAs suppress Mg^2+^ lethality *in vivo*

Finally, we sought to identify the source of Mg^2+^ growth inhibition in the ΔRLPAs Δ*cwlO* background. Because LytE is essential in the absence of CwlO, we hypothesized that the sensitivity of the Δ40 Δ*cwlO* strain to Mg^2+^ (and, by extension, the sensitivity of the ΔRLPAs *ΔcwlO* strain to Mg^2+^) could be explained by Mg^2+^ inhibition of LytE. To investigate this, we first characterized the response of Δ*cwlO* cells to the removal of LytE. We constructed an otherwise wildtype strain with *cwlO* knocked out and *lytE* under inducible control and monitored its growth by time-lapse phase-contrast microscopy. When *lytE* was induced, cell growth was normal (Movie S1). When *lytE* induction was removed, cell growth initially slowed, followed by a period of ‘stuttery’ growth, where elongating cells intermittently shrank while showing accompanying fluctuations in their phase contrast signal (Movie S2). Ultimately, cells lysed about 1-2 doubling times after the removal of *lytE* induction, as previously observed (10, 12). Next, we performed the same imaging in the ΔRLPAs *ΔcwlO* strain after the addition of 10 mM MgCl_2_ and observed the same ‘stuttery’ phenotype, suggesting that LytE function might be inhibited by Mg^2+^ (Movie S4). Without the addition of Mg^2+^, cell growth of the ΔRLPAs *ΔcwlO* strain was normal (Movie S3). In WT cells or *ΔcwlO* cells, the presence of Mg^2+^ has no effect on cell viability or growth – growth is only inhibited in the absence of the RLPAs. Thus, the RLPAs appear to allow LytE to maintain its activity in the presence of Mg^2+^. This ΔRPLAs ΔcwlO strain additionally had a similar environmental stress response profile as the Δ40 ΔcwlO strain (Figure 7A).

**Figure 7:**
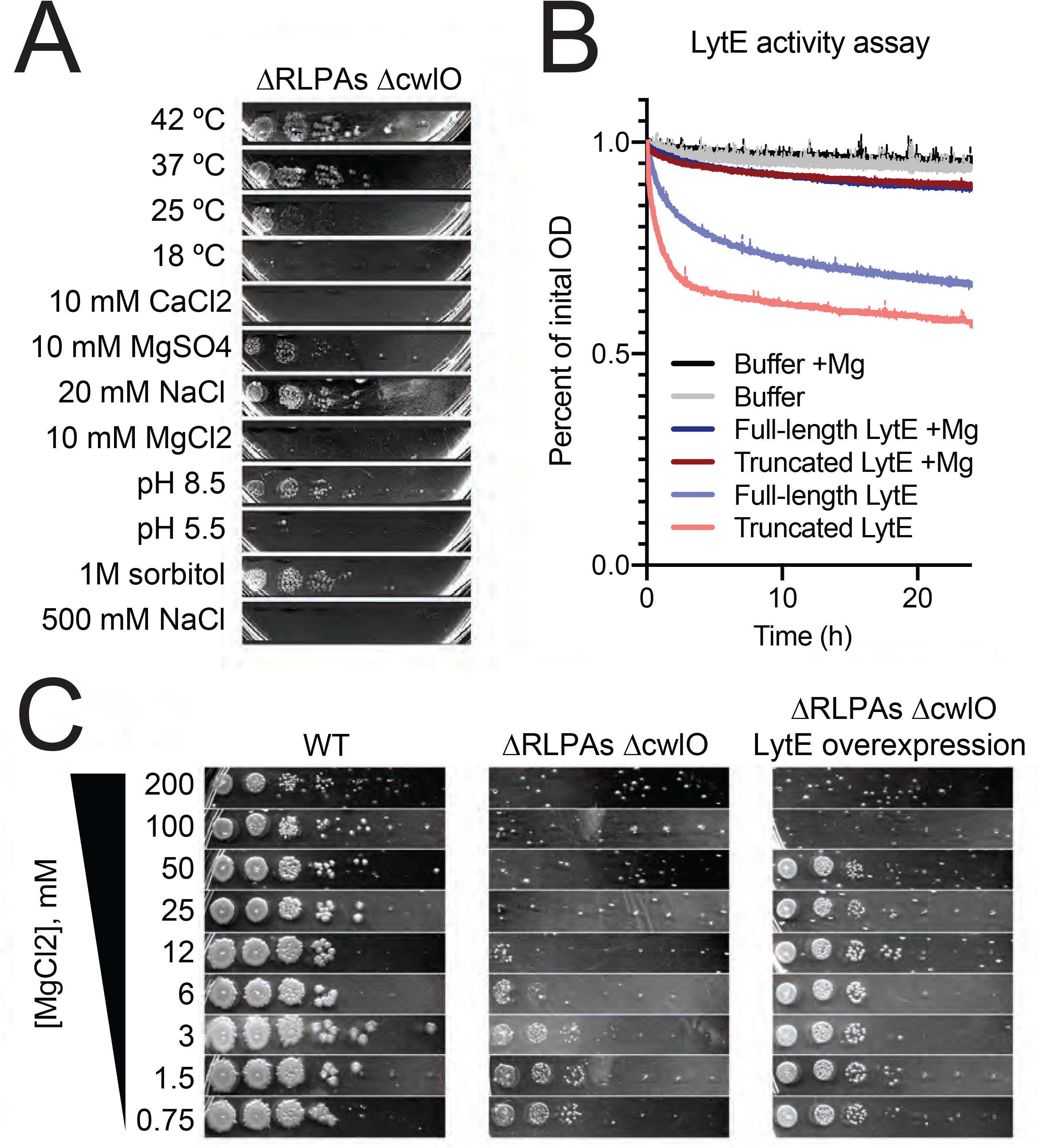
Three uncharacterized RlpA-like proteins stimulate LytE activity in the presence of divalent cations. **A: The removal of three RlpA-like proteins makes Δ*cwlO* cells stress-sensitive.** Spot dilution assays were performed as in Figure 4. Δ*yabE* Δ*yocH* Δ*ydjM* (ΔRLPAs) Δ*cwlO* showed the same stress sensitivity profile as Δ40 Δ*cwlO*, except that 10 mM MgSO_4_ and 25°C only partially inhibited growth. Strain used: bSW490, Δ*cwlO* Δ*yabE* Δ*yocH* Δ*ydjM*. **B: Mg^2+^ directly inhibits LytE activity.** LytE was expressed in E. coli and purified both with and without its N-terminal LysM domains and incubated with purified cell walls in 150 µL of 50 mM HEPES pH 7, 2 mM DTT, 500 mM NaCl with 0.5% (w/v) Pluronic F-108 in a 96-well plate. OD_600_ was measured every 2 min using a plate reader. The addition of 25 mM MgCl_2_ almost completely inhibited the activity of LytE. **C: LytE overexpression rescues Mg^2+^ sensitivity in the ΔRLPAs Δ*cwlO* background.** Spot dilutions were performed as in Figure 5A, with the indicated concentration of MgCl_2_ and the addition of 1 mM IPTG to drive LytE overexpression. Strains used: PY79, WT; (ΔRLPAs) Δ*cwlO*, bSW490, Δ*cwlO* Δ*yabE* Δ*yocH* Δ*ydjM*; (ΔRLPAs) Δ*cwlO* lytE overexpression, bSW519, Δ*cwlO* Δ*yabE* Δ*yocH* Δ*ydjM amyE::pHyperSpank-lytE*.

To test whether LytE activity is directly inhibited by Mg^2+^, we overexpressed and purified both full-length LytE and a truncated LytE protein with only its catalytic domain. *In vitro* activity assays with and without the addition of Mg^2+^ showed that indeed LytE activity is inhibited by Mg^2+^ (Figure 7B). Additionally, we reasoned that if the Mg^2+^-sensitivity phenotype was due to direct inhibition of LytE by Mg^2+^, increasing the levels of LytE should protect cells from death by increasing the total amount of LytE activity. Indeed, overexpression of LytE allowed the ΔRLPAs strain to survive in the presence of higher levels of Mg^2+^, although 100 mM MgCl_2_ still inhibited growth (Figure 7C).

Thus, we conclude that LytE activity is inhibited by Mg^2+^ both *in vivo* and *in vitro*. Furthermore, our data indicates that the RPLAs allow LytE to maintain normal function in the presence of Mg^2+^, though the specific mechanism is unclear. Whether the RLPAs act directly or indirectly on LytE remains to be determined, but we anticipate that the RLPAs interact with and activate LytE similar to what has been observed for the *Mycobacterium smegatis* hydrolases RipA and RpfB: RipA’s C-terminus (containing a NLPC/P60 domain like LytE) interacts with RpfB’s RlpA-like LTG domain (36), and RipA and RpfB have synergistic activity *in vitro* (37). By analogy, LytE’s catalytic NLPC/P60 domain may interact with the RlpA-like domains in YabE, YdjM, and YocH, leading to increased LytE activity, allowing LytE to continue to function in the presence of Mg^2+^. The ΔRLPAs Δ*cwlO* strain also has increased sensitivity to ionic stress and low temperatures, suggesting RLPAs might stimulate LytE activity under those conditions as well.

## DISCUSSION

Bacterial cell growth requires the action of PG hydrolases, but previous *in vivo* hydrolase studies have been impeded by their diversity and redundancy. We constructed and validated a *B. subtilis* strain lacking all hydrolases potentially involved in cell growth besides LytE and CwlO. These deletions constitute 40 genes in total, representing 10% of secreted proteins and 1% of all genes. The resulting Δ40 strain enables the investigation of given hydrolases and the cellular contexts in which they function, and in this work, allowed several new discoveries regarding their sufficiency, regulation, and genetic interplay.

First, we found that the Δ40 strain is viable. This demonstrates that LytE and CwlO alone can function to expand the cell wall to allow cell growth. Furthermore, as single knockouts of LytE and CwlO in the Δ40 strain are viable and allow growth (albeit at a somewhat reduced rates with some shape defects), this demonstrates *B. subtilis* requires only one of these two hydrolases to grow.

Our minimal hydrolase strain allowed us to show that RlpA-like lytic transglycosylases enhance LytE activity *in vivo* and that this enhancement can be important for growth under conditions where LytE activity is inhibited, including the presence of divalent cations, ionic stress, and cold. Although the mechanism for LytE enhancement is unclear, we hypothesize that RlpAs stimulate LytE activity via a direct interaction, as has been observed for similar proteins in *M. smegmatis* (37). Synthetic lethal or synthetic sick interactions are straightforward to identify and characterize in the Δ40 strain, giving a useful tool to interrogate genetic relationships between different hydrolases or between hydrolases and other genes of interest – such as those involved in cell wall synthesis.

Surprisingly, the growth rate of the Δ40 strain is only slightly impaired under standard lab conditions. What, then, is the function of these 40 hydrolases, and why does *B. subtilis* encode so many of them? This multitude of hydrolases likely arises from the fact that hydrolases are involved in other processes aside from cell growth such as sporulation (4) and cell motility (38). Additionally, some hydrolases might be only be needed under nutrient conditions not tested here, such as during phosphate limitation where teichoic acids are not produced, where cells may require hydrolases that are not regulated by teichoic acids (39–41). Finally, these other hydrolases may be important during non-exponential growth states such as during stationary phase, where the recycling of cell wall turnover products, lacking in the Δ40 strain, reduces cell lysis (42). Thus, a broader screen of the sensitivity of the Δ40 strain in different nutrient and environmental conditions will allow the determination of which hydrolases are useful for which conditions.

In summary, the Δ40 minimal hydrolase strain provides a powerful experimental background to investigate the function, regulation, and interplay of hydrolases, improving our understanding of precisely how these enzymes conduct their cellular tasks. In the future, individual hydrolases can be reintroduced into the Δ40 strain to investigate their specific activities in the absence of confounding contributions from the other 39 genes. Using the Δ40 strain, PG profiling can determine the biochemical activity of hydrolases. Uncovering synthetic genetic interactions between hydrolases and other genes of interest – now easy to do for all 40 hydrolases at once – will allow us to flesh out our understanding of bacterial cell growth. Understanding the function of cell wall hydrolases is essential for a complete understanding of how bacteria grow, and the Δ40 strain will allow rapid progress to this end.

## ACKNOWLEDGEMENTS

We would like to thank Carl Wivagg, Alex Bisson, Matthew Holmes, Ferran Garcia-Pichel, and Susanne Neuer for helpful advice and discussions, and Georgia Squyres for both helpful advice, discussions, and reading of the manuscript. We thank David Rudner and Yannik Brunet for plasmids. This work was funded by National Institutes of Health Grants DP2AI117923-01 to ECG, as well as support from the Volkswagen Foundation. This work was supported by the NSF-Simons Center for Mathematical and Statistical Analysis of Biology at Harvard (1764269) and the Harvard Quantitative Biology Initiative. Some work was performed at the Center for Nanoscale Systems at Harvard University, supported by NSF ECS-0335765. HPLC-MS work was performed by the Harvard Center for Mass Spectrometry Core Facility, and sequencing was performed by the Bauer Core Facility at Harvard University. We also acknowledge the support of the Wellcome Trust (212197/Z/19/Z).

## METHODS

### Strains, media, and growth conditions

Glycerol stocks stored at -80°C were struck onto LB agar plates. For strain bSW61 (*lytE::pSpac-lytE, ΔcwlO*), these plates were additionally top spread with 1 mM IPTG. After incubation overnight at 37°C, colonies were inoculated into 1 mL media (the specific media used depended on the experiment, see figure legends for details) and grown on a roller at 37°C until they reached mid-exponential-phase growth (OD_600_ ∼0.2). Cells were diluted 1:10 in prewarmed media and again grown until mid-exponential phase; this process was repeated until the start of the experiment. Alternately, a 1:10 dilution series of cells were grown overnight in media on a roller at 25 ° C. The next day, the culture whose OD_600_ was nearest to 0.2 was diluted 1:10 and grown in media at 37°C as above. S7_50_AA indicates S7_50_ media with added amino acids as in (43). CH indicates casein hydrolysate media, as in (44). LB indicates Luria Broth (Lennox) media for liquid media experiments and Luria Broth (Miller) for solid media (plates).

### Strain construction

The wild type strain for this work was *B. subtilis* PY79. Strains used in this study are listed in Table S4. Constructs were created using Gibson assembly of PCR products. Linear Gibson assembly products were transformed into competent *B. subtilis*. Transformants were selected on LB plates containing the appropriate antibiotic. The resulting strains were verified by PCR. Constructs used in this study, as well as any plasmids used to create each construct, are listed in Table S4. Primers, along with strain construction details, are listed in Table S5. Resistance cassettes and promoters were amplified from purified plasmids (listed in Table S4), all other fragments were amplified from WT gDNA.

To combine knockouts, the parent strain was transformed with PCR product containing the locus (homology arms + resistance cassette) or gDNA as indicated. All resistance cassettes used have loxP sites flanking the cassette, allowing Cre-based loop out using plasmid pDR244 (a gift from David Rudner) of the cassette to yield a markerless knockout. Removal of the plasmid was accomplished by shifting streaks to 42°C where it cannot be replicated due to a temperature-sensitive origin. Successful loop outs were confirmed via loss of antibiotic resistance.

### PHMMER search

We used pfamscan version 1.6 to search the *B. subtilis* 168 and PY79 proteomes for all pfam domains using default parameters: e-value: 0.01, significance E-values [hit]: 0.03, significance bit scores [sequence]: 25, significance bit scores [hit]: 22. We then filtered the list for domains of interest using the list of domains in Table S2, and identified putative membrane-bound/cytoplasmic proteins using UniProt (45).

### PG purification, HPLC conditions, and MS data analysis

PG purification was conducted as in (46), with an HF treatment step instead of HCl to remove teichoic acids and the addition of a protein digestion step. Cells were grown in a baffled flask to an OD_600_ of ∼0.5 in 50 mL of CH media. Cells were mixed 50/50 with 50 mL of boiling 10% SDS and boiled for 15 min in a water bath, then pelleted at 5000x g and washed 5x with ddH2O. Cells were then resuspended in 2 mL DNAse/RNase buffer (10 mM Tris pH 7.5, 2.5 mM MgCl_2_, 0.5 mM CaCl_2_) with 20 µL DNAse I and 20 µL RNAse A, then incubated overnight at 37°C and washed 3x with ddH2O to remove nucleic acids. Next, cells were resuspended in 2 mL Proteinase K buffer (10 mM Tris pH 7.5, 1 mM CaCl_2_) with 20 µL Proteinase K, incubated overnight at 45^0^C, and washed 3x with ddH2O to remove proteins. Next, cell walls were treated with 48% (v/v) hydrofluoric acid on ice for 24H, then washed twice with 100 mM Tris pH 8 and 4 times with ddH2O. Then, the PG was resuspended in 12.5 mM NaHPO_4_ pH 5.5 with 5000 units of mutanolysin and digested overnight (16h) at 37°C on a roller to yield soluble muropeptides. Undigested material was pelleted by spinning at 16000x g for 5 mins and the supernatant was transferred to a new tube. Soluble muropeptides were reduced with sodium borohydride (1 mg/mL) for 30 mins and the reaction was stopped by adding 10 µL 30% phosphoric acid. The pH was adjusted to 4-6 using NaOH, and the reduced soluble muropeptides were characterized by high-resolution LC-MS operating in both positive and negative mode. Soluble reduced muropeptides were separated on a Waters column with the following method: column temperature 52°C, flow rate 0.5 mL/min, linear gradient of solvent A (0.1% (w/v) formate) to 10% solvent B (acetonitrile + 0.1% (w/v) formate) over 80 min.

Feature detection was performed on the raw MS data using Dinosaur (47). Feature detection was done separately on both the positive and negative mode scans with default parameters. Feature data were analyzed using a custom MATLAB program, available at https://bitbucket.org/garnerlab/wilson_40_2020/. We first filtered feature data for charge < 3. Next, we filtered for the top 10 features present during each scan. For each of these features, theoretical m/z values were compared with observed m/z with a cutoff of 10 ppm. We required that a compound be present on both the positive and negative scans, and consolidated features matching the same compound within a retention time within 1 min. Finally, we filtered out compounds corresponding to in-source decay (loss of glucosamine without a change in retention time), and compounds present at less than 0.1% of all muropeptides. Retention times shown in Table S3 were analyzed manually.

### Growth rates

Cells were grown to an OD_600_ of ∼0.3-0.5 on a roller drum at 37°C and diluted to an OD_600_ of ∼0.05 in baffled flasks in a water bath shaker at 37°C. Samples were withdrawn at 5 min intervals and OD_600_ was measured in a plastic cuvette using a Biowave Cell Density Meter CO8000. T vs. OD_600_ curves were fit to a single exponential (OD_600_ = Ae^BT^) to extract a growth rate (B).

### Autolysis rates

Cells were grown to an OD_600_ of ∼0.3-0.5 on a roller drum at 37°C and diluted to an OD_600_ of ∼0.025 into prewarmed CH in baffled flasks at 37°C. Once cells reached an OD_600_ of 0.5, sodium azide (75 mM final) or ampicillin (100 µg/mL final) was added to part of culture and transferred to a prewarmed 96 well plate. OD_600_ was measured every 2 minutes for 24H at 37°C using a BioTek Epoch 2 Microplate Spectrophotometer. The plate was shaken at maximum RPM in between measurements.

### Sporulation efficiency

Sporulation was induced by resuspension according to (44). Cells were grown to an OD_600_ of ∼0.3-0.5 in CH media, pelleted, and resuspended in resuspension medium. Sporulation efficiency was assessed by measuring the number of heat-resistant CFUs per mL of culture after 36H. The cultures were heated to 80°C for 20 mins, then plated. CFU counts were then done after 24H of incubation at 37°C.

### Turnover rates

Turnover rates were measured as in (43) with some modifications; the method is summarized below. Cells were grown in S7_50_AA to an OD_600_ of ∼0.3-0.5 on a roller drum at 37°C and diluted to an OD_600_ of ∼0.05 in 3 mL of prewarmed S7_50_AA containing 1 uCi of ^3^H-N-acytylglucosamine [6-^3^H] (specific activity: 20 Ci/mmol, American Radiolabeled Chemicals, Inc., St. Louis, MI, USA) in 25mm wide test tubes in a water bath shaker at 37°C. Cells were labeled for 3 generations (until OD_600_ ∼0.4), then filtered, washed twice with prewarmed S7_50_AA, and resuspended in 25 mL of prewarmed S7_50_AA. Samples were withdrawn at 5 min intervals and OD_600_ was measured in a plastic cuvette using a Biowave Cell Density Meter CO8000. Samples were mixed 50:50 with ice cold 10% (v/v) TCA + 20 mM unlabeled GlcNAc, incubated on ice for 10 mins, then filtered and washed. Filters were dried, resuspended in Ultima Gold LSC cocktail (PerkinElmer, Waltham, MA, USA) and radioactivity was measured using a scintillation counter (Tri-Carb 2100 TR, PerkinElmer). Decays/min vs. OD_600_ plots were fit to a single exponential (DPM = Ae^BT^) to extract a turnover rate (B).

### Cell dimensions

Cells were grown to an OD_600_ of ∼0.3-0.5 in a water bath shaker at 37°C. 1 mL of culture was stained with FM 5-95 and concentrated to 100 µL by centrifugation at 2000x g and resuspension. 5 µL of concentrated cells were spotted under 2% (w/v) agarose pads in CH containing 0.5 ug/mL FM 5-95. Images were collected on a Nikon Ti-E microscope using a Nikon CFI Plan Apo DM Lambda 100X Oil objective, 1.45 NA, phase ring Ph3 using an ORCA-Flash4.0 V2 sCMOS camera. Analysis was performed using Morphometrics v1.1 (48). Zero length or width cells were discarded, as well as any cells with width greater than length. Outliers were removed using Graphpad Prism ROUT with default parameters (1%).

### Chain length

Cells were grown to an OD_600_ of ∼0.3-0.5 on a roller drum at 37°C and diluted to an OD_600_ of ∼0.025 into prewarmed CH in baffled flasks at 37°C. Once cells reached an OD_600_ of 0.5, 1-5 µl of culture was spotted under a prewarmed 2.5% (w/v) agarose pad in CH. A 10×10 image tile series was collected (∼1.5 mm square). A custom MATLAB program was used to register and stitch the images together, and then chain length was measured manually with the assistance of a custom MATLAB program.

### Electron microscopy and cell wall thickness measurements

Electron microscopy was performed as in (49). Briefly, exponentially growing cells were fixed in 100 mM MOPS buffer pH 7 containing 2% (w/v) paraformaldayde, 2.5% (w/v) gluteraldehyde, and 1% (v/v) dimethyl sulfoxide overnight at 4°C, washed, stained with 2% (w/v) osmium tetroxide in 100 mM MOPS for 1 hr, washed, and stained overnight with 2% (w/v) uranyl acetate. The cells were then dehydrated and embedded in Embed 812 resin.

Serial ultrathin sections (80 nm) were cut with a Diatome diamond knife (EMS, PA) on a Leica Ultracut UCT (Leica Microsystems, Germany) and collected on 200-mesh thin-bar formvar carbon grids. Sections were imaged on a Hitatchi HT7800 transmission electron microscope.

Images collected were segmented (inner cell wall, outer cell wall) using DeepCell (50), and cell wall thickness was measured using a custom Matlab program available at https://bitbucket.org/garnerlab/wilson_40_2020/. Briefly, the distance between the inner and outer cell wall was measured every 10 nm along a user-defined line, and the mean of that measurement was taken to be the cell’s cell wall thickness.

### AFM imaging and cell wall thickness measurement

*B. subtilis* cells at mid-exponential phase were boiled rapidly to kill bacterial cells & inactivate any potential hydrolase activity. Cells were broken by French Press and FastPrep, then suspended in 5% (w/v) SDS and boiled for 25 min, and sacculi collected by centrifugation at 20,000 g for 3 mins. The resulting pellets were washed with distilled water to remove all traces of SDS, then re-suspended in Tris-HCl (50 mM, pH7) containing 2 mg/ml pronase and incubated at 60°C for 90 mins. The resulting sacculi were then re-suspended in LC-MS Chromasolv water for storage at -20°C.

Freshly cleaved mica discs were incubated with Cell-Tak™, (285 ml 100 mM NaHCO_3_ (pH 8) then 10 µl of Cell-Tak (Corning, 5% (w/v) in acetic acid) and 5 µl of 1 M NaOH, covered and left for 20 minutes then washed five times with HPLC grade water) to ensure attachment of sacculi on the glass surface. Sacculi were diluted in HPLC-grade water to appropriate concentration and dried onto mica using N_2_. These were further washed and dried with N_2_ again to remove any unattached sample.

All AFM data was taken on a JPK Nanowizard III in QI (quantitative imaging) mode. Samples were imaged in HPLC Grade water using a FastScanD cantilever (Bruker, Santa Barbara), nominal spring constant 0.25 N/m with a 256 × 256-pixel scan region, driven at ∼167 Hz with a typical Z length of ∼ 300 nm using peak interaction forces of 2-3 nN. Images were flattened to median of differences and first order planefit using Gwyddion.

### Spot dilution assay

Cells were grown to an OD_600_ of 0.5 and diluted 1:10 into 100 µL of LB media in a 96 well plate. A 1:10 serial dilution series was made, and 3 µL of each dilution was spotted onto the plate using a multichannel pipettor. The plates were allowed to dry and incubated in at 37°C or 42°C as indicated for 18h. Plates incubated at 25°C or 18°C were left for additional time (24h and 48h, respectively). Plates were photographed using a Canon SC1011 scanner with the lid open.

For the colony morphology assay in Figure 1, this protocol was followed except that a colony of cells of each strain were simply resuspended in 100 µL of media using a toothpick (omitting the broth culture step).

### LytE purification

His-SUMO-tagged full length LytE missing the signal peptide (26–355) and His-SUMO-tagged truncated LytE missing its 3x LysM domains (185–255) were overexpressed and purified under denaturing conditions from E. coli BL21. 3L of LB Amp culture was grown to an OD_600_ of 0.7 and overexpression was induced with 1 mM IPTG. Cultures were induced for 6H, then harvested by centrifugation. Pellets were frozen at -80°C for storage. For purification, pellets were thawed and resuspended in lysis buffer (100 mM sodium phosphate, 10 mM Tris, 10 mM imidazole, 1% (v/v) Trition, 8M Urea, pH 8). Cells were lysed by sonication, and cell debris was removed by centrifugation. A His column was equilibrated in lysis buffer (3 mL bed volume). Clarified lysate was passed through the His column. The bound protein was washed once with 50 mM HEPES, 150 mM NaCl, 20 mM imidazole, 1% (v/v) Triton, 8M Urea, pH 8. Proteins were renatured on the column in 50 mM HEPES, pH 8 with 1% (v/v) Triton and 20 mM imidazole with a steady reduction of urea concentration (8M, 6M, 4M, 2M, 1M, 0M) and increasing NaCl concentration (150 mM, 237.5 mM, 325 mM, 412.5 mM, 456.25 mM, 500 mM). Refolded proteins were eluted from the column by increasing the imidazole concentration to 250 mM. DTT was added to 1 mM to all fractions. Fractions containing the target protein were pooled and dialyzed overnight at 4°C in cleavage buffer (50 mM HEPES pH 8, 500 mM NaCl, 2 mM DTT) with the addition of purified Ulp1 to cleave the 6His-SUMO tag. Buffer was exchanged once and further dialyzed for several hours. A new column was equilibrated in cleavage buffer without DTT and the pooled, cleaved protein was run through the column to remove the 6His-SUMO tag. Fractions contained cleaved protein were pooled and concentrated to a volume of 2 mL, then stored in dialysis in cleavage buffer. Activity tests were performed using purified PG in cleavage buffer plus 0.5% (w/v) Pluronic F-108 and PG from Sigma to an OD_600_ of 0.25 at 37°C. OD_600_ was measured every 2 minutes for 24H using a BioTek Epoch 2 Microplate Spectrophotometer. The plate was shaken at maximum RPM in between measurements.

### Data availability

All custom software used in this work is available at https://bitbucket.org/garnerlab/wilson_40_2020/. Raw HPLC-MS data for PG profiling experiments available from ftp://massive.ucsd.edu/MSV000086886 (doi:10.25345/C5R21D). Raw and error corrected sequencing reads for whole genome sequencing available at https://www.ncbi.nlm.nih.gov/Traces/study/?acc=PRJNA702153 (BioProject PRJNA702153).

