## Supplemental text for "An Exhaustive Multiple Knockout Approach to Understanding Cell Wall Hydrolase Function in *Bacillus subtilis*"

**Supplemental Table 1: 8 SNPs are present in the Δ40 strain, all in pathways unrelated to exponential phase cell wall synthesis.**

| Change | Codon Change | AA Change | Gene Name | CDS Name | CDS Position | Coverage | Protein Effect | Locus |
|---|---|---|---|---|---|---|---|---|
| T -> A | TCA -> ACA | S -> T | mfd | transcription-repair coupling factor CDS | 1216 | 95 | Substitution | BSU00550 |
| T -> G | CAA -> CCA | Q -> P | spsB | spore coat polysaccharide biosynthesis protein SpsB CDS | 1160 | 61 | Substitution | BSU37900 |
| T -> C | ATC -> ACC | I -> T | pucL | uric acid degradation bifunctional protein PucL CDS | 194 | 46 | Substitution | BSU32450 |
| G -> A | CCA -> CTA | P -> L | panD | aspartate 1-decarboxylase CDS | 365 | 31 | Substitution | BSU22410 |
| C -> A | TGG -> TGT | W -> C | dhbF | non-ribosomal peptide synthetase CDS | 3732 | 9 | Substitution | BSU31960 |
| T -> C | GCA -> GCG | | opuBA | choline transport ATP-binding protein OpuBA CDS | 447 | 58 | None | BSU33730 |
| G -> T | CGG -> CGT | | ykvU | sporulation protein YkvU CDS | 603 | 37 | None | BSU13830 |
| G -> A | GCC -> GCT | | czcD | H+/K+ antiporter CDS | 504 | 30 | None | BSU26650 |

Whole-genome sequencing (Illumina, paired-end) was performed on the Δ40 strain. Libraries were prepared using an Illumina Nextera XT kit. QC was performed using qPCR and TapeStation, and sequencing was performed using an Illumina NextSeq 500. 8 SNPs were detected at >8x coverage at 100% variant frequency, listed here. Strains used: bSW431, Δ40.

**Supplemental Table 2: List of PFAM domains with cell wall hydrolase activity included in our search.**

| Pfam accession | Domain name | Example protein | Reference |
|---|---|---|---|
| PF01510 | Amidase_2 | XlyA | (1) |
| PF01520 | Amidase_3 | LytC | (1) |
| PF12671 | Amidase_6 | YhbB | (3) |
| PF07454 | SpoIIP | SpoIIP | (4) |
| PF00144 | Beta-lactamase | PbpX | (5) |
| PF01915 | Glyco_hydro_3_C | NagZ | (2) |
| PF00704 | Glyco_hydro_18 | SleL | (6) |
| PF01832 | Glucosaminidase | LytD | (1) |
| PF06725 | 3D | YocH | (7) |
| PF07486 | Hydrolase_2 | CwlJ | (1) |
| PF01464 | SLT | CwlP | (1) |
| PF03330 | DPBB_1 | YdjM | (9) |
| PF08486 | SpoIID | SpoIID | (10) |
| PF00877 | NLPC_P60 | LytE | (1) |
| PF00246 | Peptidase_M14 | YqgT | (2) |
| PF01551 | Peptidase_M23 | LytH | (2) |
| PF05708 | Peptidase_C92 | YycO | (2) |
| PF13539 | Peptidase_M15_4 | CwlK | (1) |
| PF05382 | Amidase_5 | | (2) |
| PF00933 | Glyco_hydro_3 | NagZ | (2) |
| PF00062 | Lys | | (2) |
| PF00959 | Phage_lysozyme | | (2) |
| PF01183 | Glyco_hydro_25 | | (2) |
| PF05257 | CHAP | CwlO | (2) |
| PF00877 | NLPC_P60 | LytE | (2) |
| PF03411 | Peptidase_M74 | | (2) |
| PF14718 | SLT_L | EcSlt70 | (8) |
| PF03562 | MltA | EcMltA | (8) |
| PF13406 | SLT_2 | EcMltB | (8) |
| PF11873 | Mltc_N | EcMltC | (8) |
| PF11741 | AMIN | EcAmiC | (8) |
| PF01427 | Peptidase_M15 | CwlK | (2) |
| PF13702 | Lysozyme_like | YddH | (1) |
| PF02618 | YceG | EcMltG | (11) |

Hydrolase domains were collated from 3 major reviews: (1) Smith et al. 2000, (2) Vermassen et al. 2019, and (8) van Heijenoort et al. 2011, as well as other sources as indicated.

**Supplemental Figure 1: Sequencing coverage patterns in the ∆40 strain**

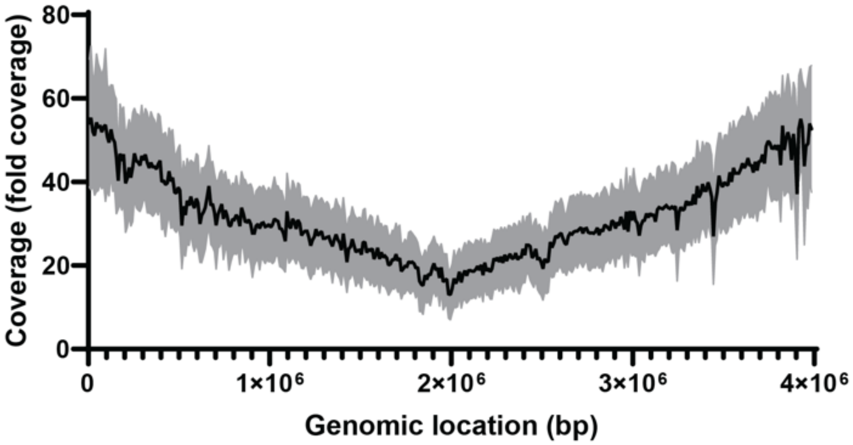


Sequencing (described in Methods and Supplemental Table 1) coverage results for the ∆40 strain. Repeat regions were excluded from coverage analysis. Mean (black line) and standard deviation (grey shading) of coverage for 1kbp regions was computed. Strains used: bSW431, ∆40.

**Supplemental Figure 2: EIC traces for several sample PG species.**

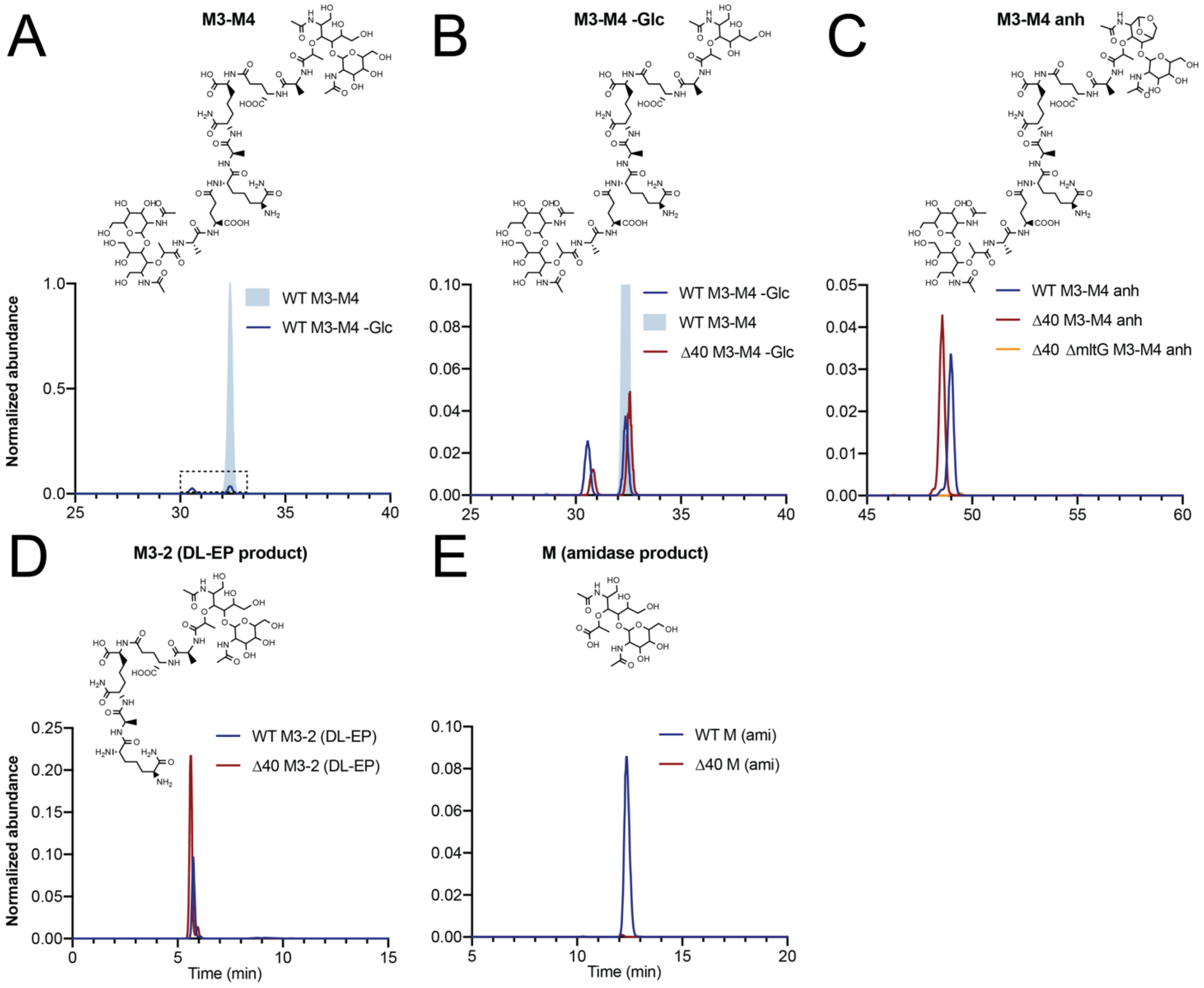

Extracted ion current (EIC) traces for several representative PG species in both WT, Δ40, and Δ40 ΔmltG strains. (A) Disaccharide tripeptide disaccharide tetrapeptide with 2 amidations (M3-M4, m/z = 896.90712, blue shaded) and disaccharide tripeptide disaccharide tetrapeptide with 2 amidations missing a glucosamine (M3-M4 -Glc, m/z = 795.36744, dark blue line) are shown. The maximum abundance of M3-M4 was used to normalize the abundance of all other species shown in all plots. (B) Zoomed view of the plot shown in A with the addition of data from the Δ40 strain. M3-M4 -Glc in the blue shaded region (same retention time as M3-M4) likely represents in-source decay of the M3-M4 PG species, while the M3-M4 -Glc with a retention time around 30 mins likely represents glucosamindase product. (C) Anhydrodisaccharide tripeptide disaccharide tetrapeptide with 2 amidations (m/z = 886.89402) abundance is shown for WT, Δ40, and Δ40 ΔmltG strains. (D) Disaccharide tripeptide dipeptide with 2 amidations (m/z = 556.76951) abundance is shown for WT and Δ40 strains. (E) Disaccharide (m/z = 499.21336) abundance is shown for WT and Δ40 strains. Strains used: PY79, WT; bSW431, Δ40; bSW537, Δ40 Δ*mltG*.

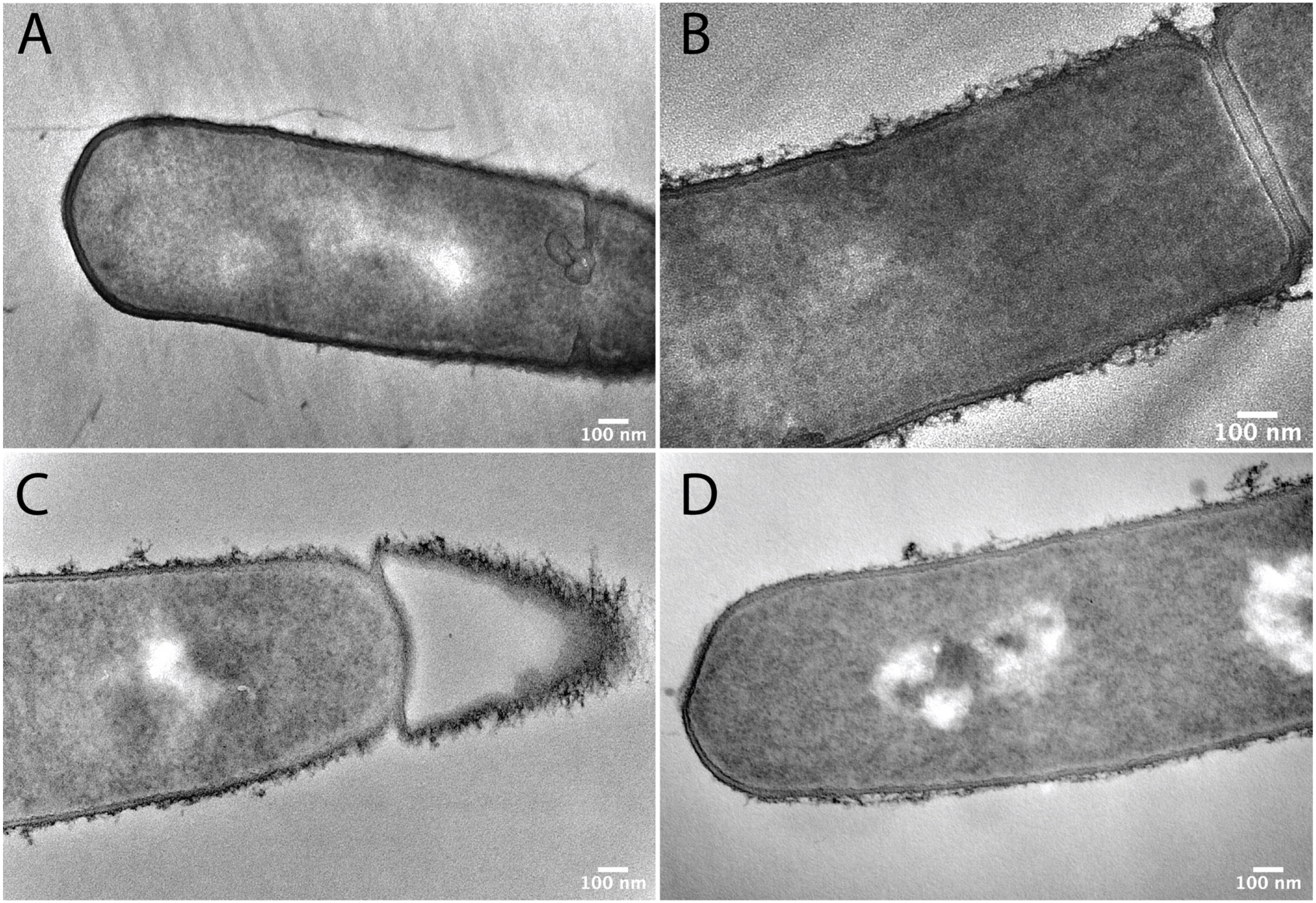
A
100 nm
B
100 nm
C
100 nm
D
100 nm

**Supplemental Figure 3: Representative image of cells via TEM.**

**A: Whole cell image of PY79 (WT)** from the dataset analyzed in Figure 4B. Strains used: PY79, WT.

**B: Whole cell image of Δ40** from the dataset analyzed in Figure 4B. Strains used: bSW431, Δ40.

**C: Whole cell image of Δ40 ΔlytE,** prepared as in Figure 4B (details in Methods). Strains used: bSW435, Δ40 ΔlytE.

**D: Whole cell image of Δ40 ΔcwlO,** prepared as in Figure 4B (details in Methods). Strains used: bSW433, Δ40 ΔcwlO.

**Supplemental Movie 1: Growth of inducible *lytE,* Δ*cwlO* strain in the presence of inducer.** Cells were spotted under an agarose pad containing media with inducer (CH + 250 uM IPTG) and imaged using phase-contrast microscopy. Frames are 1 minute apart. Strains used: bSW61, *lytE::pSpac-lytE*.

**Supplemental Movie 2: 'Stuttery' growth before lysis of inducible *lytE,* Δ*cwlO* strain upon removal of inducer.** Cells were spotted under an agarose pad containing media without inducer (CH) and imaged using phase-contrast microscopy. Frames are 1 minute apart. Strains used: bSW61, *lytE::pSpac-lytE*.

**Supplemental Movie 3: Normal growth of WT cells before and after addition of $Mg^{2+}$.** Cells were loaded into a CellASIC BO4A plate in CH media and imaged using phase-contrast microscopy. At frame 18, media was exchanged for the same media plus 20 mM $Mg^{2+}$ (indicated by label in upper left hand corner.) Frames are 2 minutes apart. Strains used: WT, PY79.

**Supplemental Movie 4: 'Stuttery' growth of ΔRLPAs Δ*cwlO* strain only after addition of $Mg^{2+}$** Cells were loaded into a CellASIC BO4A plate in CH media and imaged using phase-contrast microscopy. At frame 18, media was exchanged for the same media plus 20 mM $Mg^{2+}$ (indicated by label in upper left hand corner.) Frames are 2 minutes apart. Strains used: bSW490, ΔRLPAs Δ*cwlO*.
